# Gluconeogenic Enzyme PCK1 Deficiency Is Critical for CHK2 O-GlcNAcylation and Hepatocellular Carcinoma Growth upon Glucose Deprivation

**DOI:** 10.1101/2020.08.06.240739

**Authors:** Jin Xiang, Chang Chen, Rui Liu, Dongmei Gou, Lei Chang, Haijun Deng, Qingzhu Gao, Wanjun Zhang, Lin Tuo, Xuanming Pan, Li Liang, Jie Xia, Luyi Huang, Ailong Huang, Kai Wang, Ni Tang

**Affiliations:** Key Laboratory of Molecular Biology for Infectious Diseases (Ministry of Education), Institute for Viral Hepatitis, Department of Infectious Diseases, The Second Affiliated Hospital, Chongqing Medical University, Chongqing, China; Institute of Life Sciences, Chongqing Medical University, Chongqing, China; State Key Laboratory of Proteomics, Beijing Proteome Research Center, National Center for Protein Sciences (Beijing), Beijing Institute of Lifeomics, Beijing, China; Sichuan Provincial People’s Hospital, Sichuan, China

## Abstract

Elevated hexosamine-biosynthesis pathway (HBP) activity and O-GlcNAcylation are emerging hallmarks of hepatocellular carcinoma (HCC). Inhibiting O-GlcNAcylation could be a promising anti-cancer strategy. Here, we investigate this possibility and demonstrate that deficiency of phosphoenolpyruvate carboxykinase 1 (PCK1), a rate-limiting enzyme in gluconeogenesis, promotes O-GlcNAcylation and hepatoma cell proliferation under low-glucose conditions. PCK1 loss results in oxaloacetate accumulation and AMPK inactivation, promoting uridine diphosphate-N-acetylglucosamine (UDP-GlcNAc) synthesis and CHK2 threonine 378 O-GlcNAcylation and counteracting its ubiquitination and degradation. O-GlcNAcylation also promotes CHK2-dependent Rb phosphorylation and HCC cell proliferation. Therefore, blocking HBP-mediated O-GlcNAcylation suppresses tumor progression in liver-specific *Pck1*-knockout mice. We reveal a link between PCK1 depletion and hyper-O-GlcNAcylation that underlies HCC oncogenesis and suggest therapeutic targets for HCC that act by inhibiting O-GlcNAcylation.

## INTRODUCTION

Gaining insight into the fundamental role of metabolic reprogramming in cancer has contributed immensely to our understanding of tumorigenesis and cancer progression ^1^. Nutrient limitations (such as glucose deprivation) in solid tumors may require cancer cells to exhibit the metabolic flexibility required to sustain proliferation and survival ^2^. Increased aerobic glycolysis (the Warburg effect) is one of main characteristics of cancer cells that supports their intensive growth and proliferation ^3^. Gluconeogenesis (the pathway opposite to glycolysis) operates during starvation and occurs mainly in the liver, and also plays key roles in metabolic reprogramming, cancer cell plasticity, and tumor growth ^4,5^. Several key gluconeogenic enzymes, such as phosphoenolpyruvate carboxykinase 1 (PCK1, also known as PEPCK-C), fructose-1,6-bisphosphatase, and glucose-6-phosphatase, were previously found to be dysregulated in several types of cancer, including hepatocellular carcinoma (HCC) ^6^.

The cytoplasmic isoform of PCK1, the first rate-limiting enzyme in hepatic gluconeogenesis, catalyzes the conversion of oxaloacetate (OAA) to phosphoenolpyruvate (PEP). The oncogenic or tumor suppressor roles of PCK1 in different types of human cancers are rather controversial. PCK1 has anti-tumorigenic effects in gluconeogenic organs (liver and kidney), but has tumor-promoting effects in cancers arising from non-gluconeogenic organs ^4^. In colon-derived tumor cells, PCK1 is hijacked to participate in truncated gluconeogenesis to meet biosynthetic and anabolic needs ^7^. However, the underlying mechanism determining its aberrant expression and altered function in multiple types of tumors remains incompletely understood.

Recent findings emphasize the role of the hexosamine-biosynthesis pathway (HBP), a sub-branch of glucose metabolism, in carcinogenesis ^8–10^. The HBP and glycolysis share the first two steps and diverge at fructose-6-phosphate (F6P). Glutamine-fructose-6-phosphate aminotransferase 1 (GFAT1), the rate-limiting enzyme of the HBP, converts F6P and glutamine to glucosamine-6-phosphate and glutamate. Uridine diphosphate N-acetylglucosamine (UDP-GlcNAc), the end products of HBP, is a donor substrate for O-linked β-N-acetylglucosamine (O-GlcNAc) modification (also known as O-GlcNAcylation) ^11,12^. O-GlcNAc transferase (OGT)-mediated protein O-GlcNAcylation is highly dependent on the intracellular concentration of the donor substrate UDP-GlcNAc, which is proposed to be a nutrient sensor that couples metabolic and signaling pathways ^13,14^. Increased glucose flux through the HBP and elevated UDP-GlcNAc contribute to hyper-O-GlcNAcylation in cancer cells ^15^. Previous data suggested that elevated O-GlcNAcylation may serve as a hallmark of cancer ^16^.

Similar to phosphorylation, O-GlcNAcylation is a dynamic post-translational modification that regulates protein subcellular localization, stability, protein–protein interactions, or enzymatic activity according to the nutrient demands of cells ^17^. OGT and O-GlcNAcase (OGA) are the only enzymes known to be responsible for adding and removing N-acetylglucosamine (GlcNAc) on serine and threonine residues of target proteins ^18^. Numerous oncogenic factors, including c-MYC, Snail, and β-catenin, are targets of O-GlcNAcylation ^19–21^. Advances in mass spectrometry (MS)-based methods will facilitate the discovery of novel O-GlcNAc-modified proteins in cancer cells. Therefore, modulating the HBP or O-GlcNAcylation (which regulate oncogenic activation in human cancers) represents a promising anti-cancer strategy that can potentially be used in combination with other treatments ^22,23^.

In this study, we explored the role of the gluconeogenic enzyme PCK1 in regulating the HBP and HCC proliferation under low-glucose conditions. We unravel a molecular mechanism responsible for enhanced UDP-GlcNAc biosynthesis and O-GlcNAcylation induced by PCK1 depletion, and delineate the functional importance of checkpoint kinase 2 (CHK2) O-GlcNAcylation in HCC tumorigenesis. Importantly, our study reveals a novel link between the gluconeogenic enzyme PCK1 and HBP-mediated O-GlcNAc modification, which suggest a therapeutic strategy for treating HCC.

## RESULTS

### PCK1 Deficiency Increases Global O-GlcNAcylation in Hepatoma Cells under Low-Glucose Conditions and Promotes HCC Proliferation

To explore the role of the gluconeogenic enzyme PCK1 in O-GlcNAcylation, we first analyzed the global O-GlcNAcylation levels of hepatoma cells in response to PCK1 modulation with various concentrations of glucose (1 to 25 mM, 12 h). In the presence of 5 mM glucose, PCK1 knockout significantly elevated the global O-GlcNAcylation levels (Fig. 1a,b), whereas overexpression of wild-type (WT) PCK1 markedly decreased the global O-GlcNAcylation levels in SK-Hep1, Huh7, and MHCC-97H cells (Fig. 1c and Extended Data Fig. 1a-c). Interestingly, the catalytically inactive G309R mutant of PCK1 ^24^ was unable to reduce the O-GlcNAcylation levels in these hepatoma cells under the same culture conditions (Fig. 1c and Extended Data Fig. 1b-c), suggesting that the enzymatic activity of PCK1 may contribute to its role in regulating cellular O-GlcNAcylation levels. Further, 3-mercaptopicolinic acid (3-MPA), a specific inhibitor of PCK1, dose-dependently increased the O-GlcNAcylation levels in PLC/PRF/5 cells (Fig. 1d), whereas dexamethasone (Dex), an activator of gluconeogenesis, lowered the O-GlcNAcylation levels in SK-Hep1 cells (Fig. 1e). In addition, this observation was confirmed via pharmacological or transcriptional inhibition of OGT and OGA (Fig. 1f-i), suggesting PCK1 inhibits OGT-mediated O-GlcNAcylation. Interestingly, PCK1 did not change the mRNA or protein expression levels of OGT, OGA, and GFAT1, the key enzymes involved in regulating O-GlcNAcylation and HBP (Extended Data Fig. 1d-m and Fig. 1d,e). In addition, our cell-proliferation assays indicated that PCK1 suppresses hepatoma cell proliferation, depending on its enzymatic activity and the cellular O-GlcNAcylation levels (Extended Data Fig. 2).

**Fig. 1.**
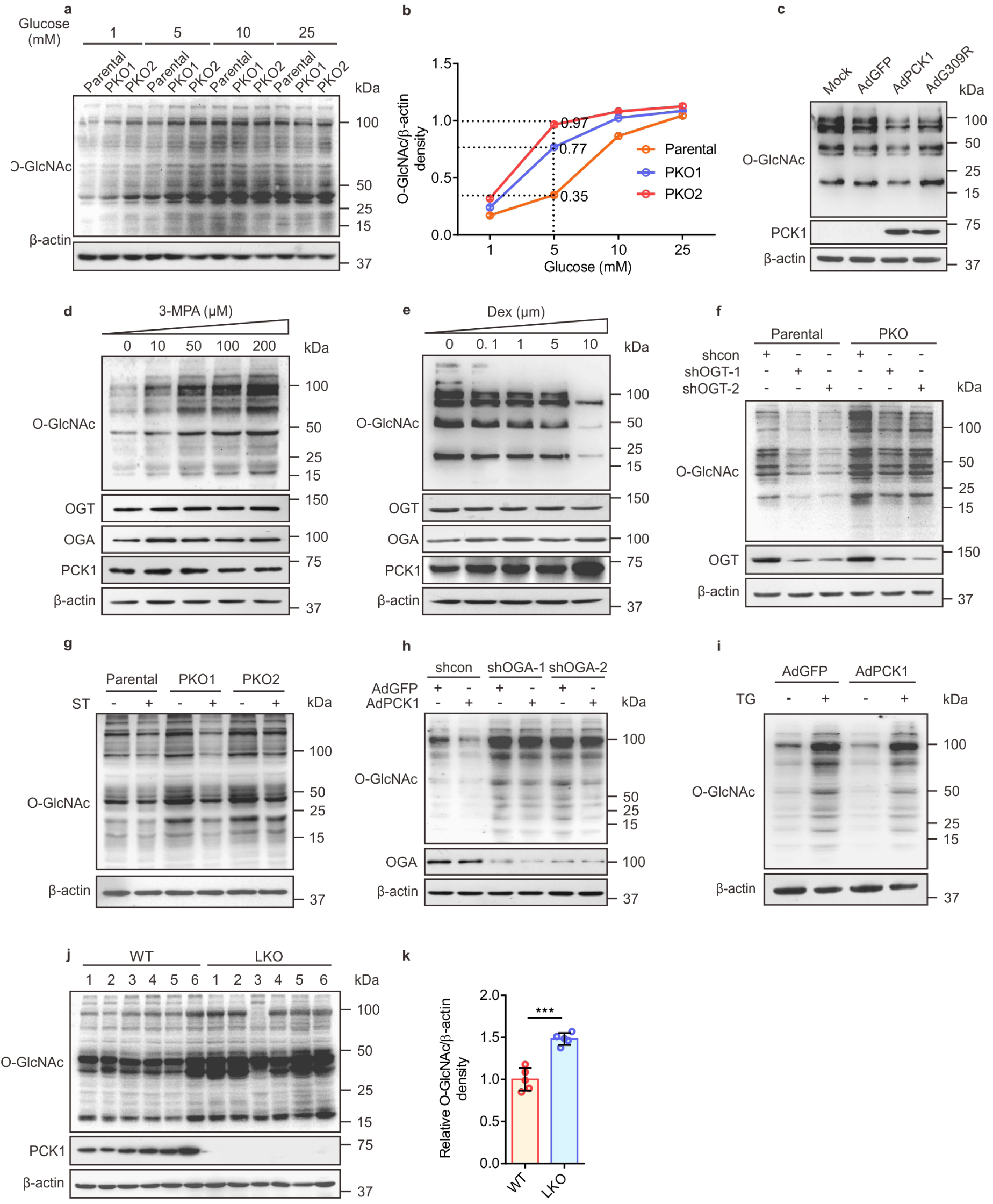
PCK1 Deficiency Enhances Protein O-GlcNAcylation. **a**,**b**, Immunoblotting analysis of global O-GlcNAcylation levels in PCK1-knockout PLC/PRF/5 cells (PKO cells) treated with medium containing different levels of glucose for 12 h (**a**). Densitometric analysis was performed with Image-pro plus software (**b**). **c**, Representative western blot analysis of the indicated proteins in SK-Hep1 cells overexpressing green fluorescent protein (GFP; control cells), WT PCK1, or an enzymatically deficient mutant (PCK1 G309R) after incubation in medium containing 5 mM glucose for 12 h. Mock-treated cells served as a blank control. **d**,**e**, Immunoblots of the indicated proteins in PLC/PRF/5 cells treated with 3-MPA for 12 h (**d**) and SK-Hep1 cells treated with Dex for 7 days (**e**). **f**,**g**, OGT and protein O-GlcNAcylation levels in PKO cells. Cells were transfected with shRNA targeting OGT mRNA or a scrambled control shRNA (shcon) for 48 h (**f**), or treated with 50 μM ST045849 (ST) for 12 h (**g**). **h**,**i**, Immunoblotting analysis of SK-Hep1 cells. PCK1-expressing cells were transfected with an OGA shRNA1/2 plasmid for 48 h (**h**), or treated with 25 μM Thiamet G (TG) for 12 h (**i**). **j**,**k**, Immunoblotting (**j**) and densitometric analysis (**k**) of liver tumors from DEN/CCl_4_-induced WT and LKO mice after fasting for 12 h. Data are represented as mean ± standard deviation (SD; n = 6 experiments). ***p < 0.001, Student’s t-test.

**Fig. 2.**
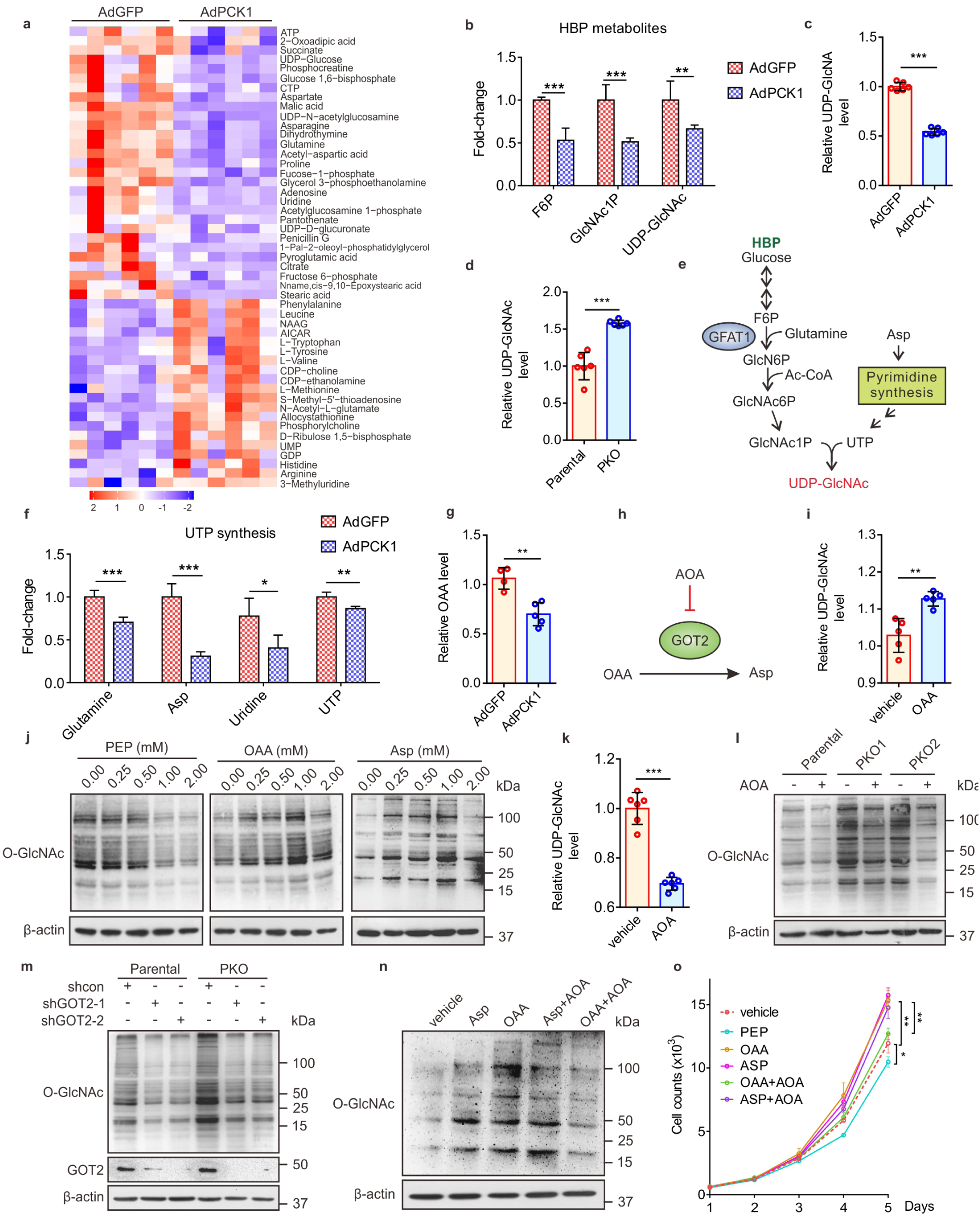
PCK1 Knockout Promotes UDP-GlcNAc Synthesis Partially Through Oxaloacetate Accumulation. **a**,**b**, Heatmap of metabolites (**a**) and fold-changes in intermediate metabolites of the HBP (**b**). **c**,**d**, Relative UDP-GlcNAc levels were measured by LC-MS in PCK1-OE SK-Hep1 cells (**c**) and PKO cells (**d**). **e**, Schematic representation of the HBP. Glucose intake feed into the HBP that produces UDP-GlcNAc. N-Acetylglucosamine-1-phosphate (GlcNAc1P) and UTP, terminal metabolites of the HBP and pyrimidine synthesis, represent the final rate-limiting steps of UDP-GlcNAc synthesis. **f**, Fold-changes in the intermediate metabolites of uridine synthesis. **g**, Relative OAA levels, as measured by LC-MS in PCK1 -OE SK-Hep1 cells. **h**, OAA is converted to Asp by the mitochondrial GOT2 enzyme, which can be prevented by the GOT2 inhibitor, AOA. **i**, Relative levels of UDP-GlcNAc, as measured by LC-MS in PKO cells treated with 1 mM OAA. **j**, Protein O-GlcNAcylation levels in SK-Hep1 cells cultured for 12 h in medium containing 5 mM glucose and PEP (left), OAA (middle), or Asp (right). **k**, Relative UDP-GlcNAc levels were measured by LC-MS in PKO cells treated with 20 μM AOA. **l**,**m**, Protein O-GlcNAcylation levels in PKO-cells treated with 20 μM AOA for 12 h (**l**) or transfected with a GOT2 shRNA1/2 plasmid for 48 h (**m**). **n**, Immunoblots of SK-Hep1 lysates treated for 12 h with OAA (1 mM), Asp (1 mM), or AOA (20 μM), as indicated. **o**, Proliferation ability of SK-Hep1 cells treated as described in (N). Data are represented mean ± SD (n ≥ 3 experiments). *p < 0.05, **p < 0.01, ***p < 0.001, Student’s t-test (two groups) or one-way analysis of variance (ANOVA) followed by Tukey’s test (more than two groups).

Additionally, we used an N-nitrosodiethylamine (DEN) and CCl_4_-induced mouse HCC model to verify the above results *in vivo*. The O-GlcNAcylation levels in the hepatic tumors of liver-specific *Pck1* knockout (LKO) mice were significantly higher than those in WT mice after 12 h of fasting (Fig. 1j,k). Furthermore, using an orthotopic HCC mouse model, we found that PCK1 overexpression or dexamethasone administration decreased the O-GlcNAcylation levels and inhibited HCC growth (Extended Data Fig. 3). Together, these data demonstrate that PCK1 decreases the global O-GlcNAcylation levels in HCC under a low-glucose condition and suppresses hepatoma cell proliferation, both *in vitro* and *in vivo*. The enzymatic activity of PCK1 is indispensable for its tumor suppressor role in HCC.

**Fig. 3.**
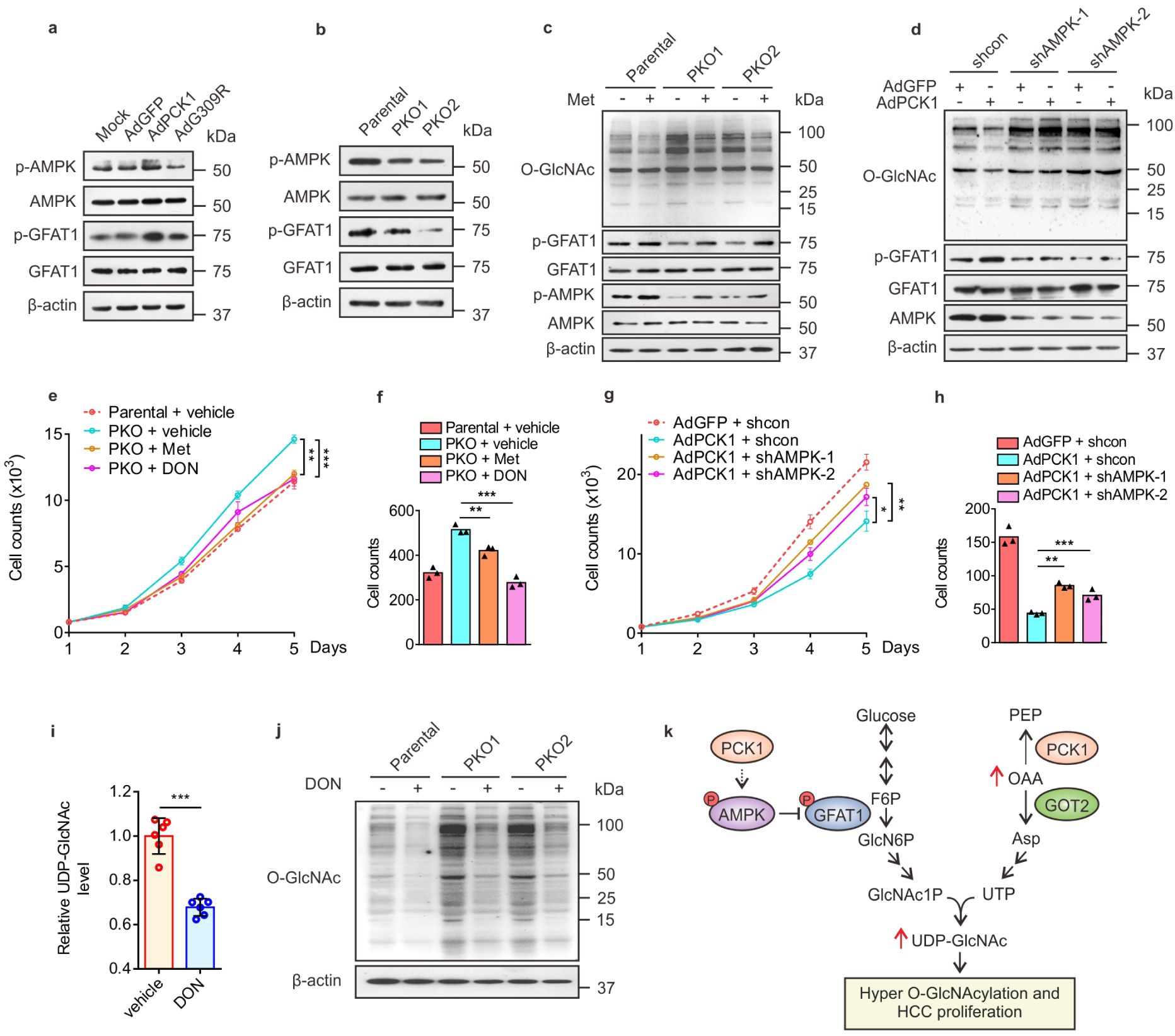
PCK1 Activates AMPK^Thr172^/GFAT1^Ser243^ Phosphorylation and Inhibits UDP-GlcNAc Biosynthesis. **a**,**b**, Representative immunoblots showing AMPK^Thr172^ and GFAT1^Ser243^ phosphorylation in PCK1-OE cells (**a**) and PKO cells (**b**). **c**,**d**, Immunoblot analysis. PKO cells were treated with metformin (Met, 2 mM) for 12 h (**c**) or transfected with an AMPK shRNA1/2 plasmid (**d**). **e**-**h**, Hepatoma cell growth curves and colony formation capacity. PKO cells and PCK1-OE cells were treated as indicated. **i**, Relative levels of UDP-GlcNAc in PKO cells treated for 24 h with the GFAT1 inhibitor DON (20 μM), as measured by LC-MS. **j**, PKO cells were treated with 20μM DON for 24 h. **k**, Working model whereby PCK1 ablation promotes UDP-GlcNAc biosynthesis, O-GlcNAcylation, and proliferation of HCC cells through increased oxaloacetate accumulation and activation of the AMPK-GFAT axis. Data are represented as mean ± SD (n ≥ 3 experiments). *p < 0.05, **p < 0.01, ***p < 0.001, as determined using Student’s t-test (two groups) or one-way ANOVA, followed by Tukey’s test (more than two groups).

### PCK1 Deficiency Promotes UDP-GlcNAc Biosynthesis via Oxaloacetate Accumulation

Next, a metabolomics assay was performed with AdPCK1- and AdGFP-infected SK-Hep1 cells to explore metabolic changes occurring after PCK1 overexpression (PCK1-OE) under a low glucose concentration (5 mM). Principal component analysis showed that PCK1 overexpression dramatically changed the intracellular metabolic profile of SK-Hep1 cells (Extended Data Fig. 4a). Levels of several metabolites of the HBP decreased after PCK1 overexpression, including fructose 6-phosphate, N-acetyl glucosamine 1-phosphate (GlcNAc-1-P), and UDP-GlcNAc (the HBP end product), as shown in Fig. 2a,b and Extended Data Fig. 4b. Our targeted liquid chromatography-tandem MS (LC-MS/MS) results showed that UDP-GlcNAc significantly decreased in PCK1-OE SK-Hep1 cells (Fig. 2c), but increased in PCK1-KO (PKO) cells (Fig. 2d), suggesting that PCK1 may negatively regulate UDP-GlcNAc biosynthesis for O-GlcNAcylation. To investigate how PCK1 modulates UDP-GlcNAc biosynthesis, we performed the pathway-enrichment analysis of metabolite profiles and found that several metabolic pathways were significantly affected, including purine and pyrimidine metabolism which required for uridine triphosphate (UTP) synthesis (Extended Data Fig. 4c). The HBP utilizes glucose, glutamine, acetyl-CoA, and UTP to produce the amino sugar UDP-GlcNAc (Fig. 2e), we assumed that PCK1 may regulate UDP-GlcNAc biosynthesis partially via UTP synthesis. Indeed, metabolomics data showed that levels of several metabolites involved in UTP synthesis including aspartate (Asp, critical metabolite in *de novo* UTP synthesis) declined upon PCK1 overexpression (Fig. 2f), besides metabolites in glycolysis and the tricarboxylic acid (TCA) cycle (Fig. 2a and Extended Data Fig. 4d–g), suggesting that PCK1 may play role in TCA cataplerosis and UTP synthesis which contribute to UDP-GlcNAc biosynthesis.

**Fig. 4.**
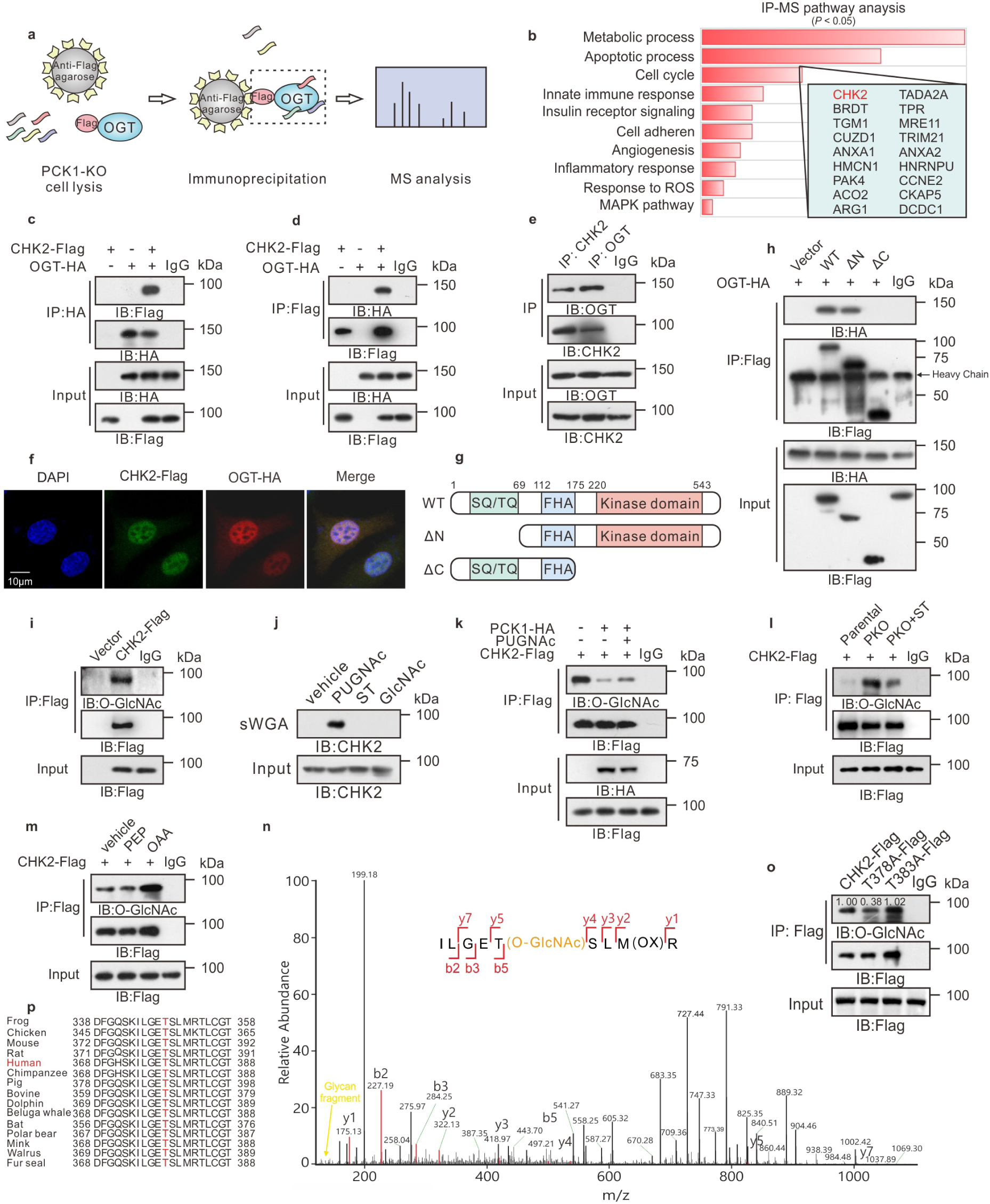
PCK1 Deficiency Promotes CHK2 O-GlcNAcylation at T378. **a**,**b**, IP-LC-MS/MS analysis of O-GlcNAc-modified proteins. (**a**) Flowchart describing the processes used for IP-LC-MS/MS analysis. (**b**) Kyoto Encyclopedia of Genes and Genomes-based analysis of significantly enriched pathways represented by proteins that bound to Flag-tagged OGT. PKO cells were transfected with OGT-Flag, and cell lysates were immunoprecipitated using anti-Flag agarose beads. **c**,**d**, Co-IP of OGT-HA and CHK2-Flag was examined using an anti-HA antibody (**c**) or an anti-Flag antibody (**d**). **e**, Co-IP of endogenous OGT and CHK2 in PKO cells. **f**, Subcellular co-localization of OGT and CHK2 in PKO cells was determined by immunofluorescence staining. **g**, Schematic representation of the CHK2 constructs. WT CHK2 contains three domains, including a SQ/TQ cluster domain, a Forkhead-associated (FHA) domain, and a kinase domain. Truncation mutants of CHK2, comprising amino acids (aa) 69–543 or 1–221, were designated as ΔN and ΔC, respectively. **h**, Interactions between OGT and full-length (aa 1–543), the ΔN truncation mutant (aa 69–543), or the ΔC truncation mutant (aa 1–221) in HEK293 cells were determined by Co-IP. **i**, CHK2 IP with anti-Flag M2 agarose beads in HEK293 cells transfected with a Flag-CHK2-expression construct or a vector control. **k**, PKO cells were treated with 50 μM PUGNAc or 50 μM ST for 24 h, incubated in 5 mM glucose, and followed by a sWGA pull-down assay. Western blot was determined by anti-CHK2. **k**-**m**, Cell lysates of PCK1-OE cells (**k**), PKO cells (**l**), or SK-Hep1 cells treated with 1mM PEP or OAA (**m**) were immunoprecipitated with anti-Flag agarose beads and immunoblotted, as indicated. **n**, LC-MS analysis of CHK2-Flag identified residue T378 as the CHK2 O-GlcNAcylation site, which corresponded to the O-GlcNAcylated CHK2 peptide ILGETSLMR. **o**, HEK293 cells were transfected with vectors encoding Flag-tagged versions of WT CHK2, T378A CHK2, or T383A CHK2. Cell lysates were purified using anti-Flag M2 agarose beads and probed with an anti-Flag or anti-O-GlcNAc antibody. Quantitative analysis of the O-GlcNAcylation/Flag (IP) band was performed using Image-pro plus software. **p**, Cross-species sequence alignment of CHK2.

PCK1 catalyzes the conversion of OAA (an intermediate of the TCA cycle) to PEP. Consistent with the results of a previous study ^7^, restoring PCK1 decreased the OAA concentration in SK-Hep1 cells (Fig. 2g). Considering that OAA is converted to Asp by the mitochondrial glutamate-oxaloacetate transaminase (GOT2, also known as aspartate aminotransferase 2; Fig. 2h), we speculated that PCK1 repression may promote UDP-GlcNAc biosynthesis through Asp conservation caused by OAA accumulation. As expected, adding OAA strengthened UDP-GlcNAc biosynthesis (Fig. 2i). Accordingly, both OAA and Asp enhanced O-GlcNAcylation in SK-Hep1 cells, while high concentrations of PEP (above 0.5 mM) showed an opposite effect (Fig. 2j). Next, we tested whether GOT2 can contribute to HBP-mediated O-GlcNAcylation in PKO cells. We found that treatment with aminooxyacetic acid (AOA), a specific inhibitor of GOT2, decreased the UDP-GlcNAc levels in PKO cells (Fig. 2k). Moreover, AOA treatment or short-hairpin RNA (shRNA)-mediated GOT2 knockdown reduced the O-GlcNAcylation levels in PKO cells (Fig. 2l,m). In addition, AOA markedly blocked O-GlcNAcylation induced by OAA treatment, but failed to moderate the effect of Asp, indicating that GOT2 plays an essential role in the metabolism of OAA converted to Asp, the HBP, and tumorigenesis (Fig. 2n,o and Extended Data Fig. 4h). Taken together, these data provide strong evidence supporting that OAA accumulation and GOT2-mediated pathway contribute to enhanced UDP-GlcNAc biosynthesis and hyper-O-GlcNAcylation in PKO cells.

### Restoration of PCK1 Suppresses O-GlcNAcylation by Activating the AMPK-GFAT1 Axis

The final rate-limiting step of UDP-GlcNAc synthesis involves UTP and GlcNAc-1-P (Fig. 2e). Our metabolomics analysis showed that both UTP and GlcNAc-1-P levels were significantly decreased in PCK1-OE cells (Fig. 2b,f). Since OAA accumulation contributes to the downstream UTP increase and GFAT1 is the rate-limiting enzyme in GlcNAc-1-P synthesis, we then explored whether GFAT1 activity also regulates UDP-GlcNAc production. Previously, we reported that PCK1 activates AMP-activated protein kinase (AMPK) upon glucose deprivation in HCC ^25^, and other groups showed that AMPK activation reduces O-GlcNAcylation through GFAT1 phosphorylation (which diminished GFAT1 activity) in endothelial cells and cardiac hypertrophy ^26,27^.

We speculated that PCK1 may also inhibit UDP-GlcNAc synthesis through the AMPK-GFAT1 axis. Thus, we tested whether PCK1 can suppress the HBP through the AMPK-GFAT1 axis under low-glucose conditions. As expected, PCK1 overexpression promoted the phosphorylation of both AMPK and GFAT1 (Fig. 3a), whereas PCK1-KO downregulated p-AMPK and p-GFAT1 production (Fig. 3b). The AMPK activator metformin partially offset hyper-O-GlcNAcylation and growth-promoting effects mediated by PCK1 depletion (Fig. 3c). However, shRNA-mediated knockdown of AMPK in PCK1-OE cells rescued the inhibitory effects of PCK1 on O-GlcNAcylation (Fig. 3d).

Furthermore, we investigated whether the inhibition of hepatoma cell proliferation in response to PCK1 depended on the AMPK-GFAT1 axis. We found that metformin suppressed PKO cell proliferation (Fig. 3e,f). In contrast, shRNA against AMPK mRNA (shAMPK) promoted PCK-OE cell proliferation (Fig. 3g,h). In addition, the GFAT1 inhibitor 6-diazo-5-oxo-l-norleucine (DON) reduced UDP-GlcNAc biosynthesis (Fig. 3i), O-GlcNAcylation levels (Fig. 3j), and PKO cell proliferation (Fig. 3e,f). These data indicate that PCK1 suppresses HBP-mediated O-GlcNAcylation and HCC proliferation partially via activation of the AMPK-GFAT1 axis. PCK1 deficiency boosts flux through the HBP and results in an increased availability of UDP-GlcNAc for O-GlcNAcylation. Therefore, both OAA accumulation and the AMPK-GFAT1 axis contributed to hyper-O-GlcNAcylation and PKO cell proliferation upon glucose deprivation (Fig. 3k).

### OGT mediates CHK2 O-GlcNAcylation in PCK1-Deficient Hepatoma Cells

To further explore how OGT-mediated protein O-GlcNAcylation facilitates hepatoma cell proliferation in PKO cells, we used immunoprecipitation coupled with tandem MS (IP-MS/MS) to screen for proteins that specifically interact with OGT (Fig. 4a). Flag-tagged OGT was transiently expressed in PKO cells, and subsequent IP-MS identified 617 candidate OGT-binding proteins (Supplemental Table 1). Gene Ontology-based biological-process analysis indicated that several proteins were involved in metabolic processes, apoptotic processes, and cell-cycle progression (Fig. 4b). We then focused on CHK2, which is required for checkpoint-mediated cell cycle arrest ^28^. Interactions between OGT and CHK2 were confirmed by co-immunoprecipitation (co-IP) experiments in HEK293 cells (Fig. 4c,d) and PKO cells (Fig. 4e). Confocal analysis also indicated that OGT and CHK2 co-localized in the nucleus (Fig. 4f). To define the precise region(s) in CHK2 required for this interaction, we expressed full-length HA-tagged OGT in combination with different Flag-tagged fragments of CHK2 in HEK293 cells (Fig. 4g). The C-terminal region of CHK2 (amino acids 69–543) containing kinase domains showed a strong interaction, whereas the N-terminal region (amino acids 1–175) did not interact with OGT (Fig. 4h).

Then, we determined whether CHK2 can be modified via O-GlcNAc. Immunoprecipitated, Flag-tagged CHK2 exhibited a strong O-GlcNAc signal in HKE293 cells (Fig. 4i). Endogenous CHK2 O-GlcNAcylation was confirmed by affinity chromatography in presence of the OGA inhibitor PUGNAc, using succinylated wheat germ agglutinin (sWGA), a modified lectin that specifically binds O-GlcNAc-containing proteins (Fig. 4j). Furthermore, PCK1-OE and the OGT inhibitor ST045849 (ST) decreased CHK2 O-GlcNAcylation (Fig. 4k,l), while PCK1-KO, the OGA inhibitor PUGNAc, or OAA treatment strengthened CHK2 O-GlcNAcylation under low-glucose conditions (Fig. 4l,m).

Next, we sought to map the O-GlcNAcylation site(s) on CHK2. Flag-tagged CHK2 was purified from PKO cells and analyzed by MS. As shown in Fig. 4n, threonine 378 (T378) was the main O-GlcNAcylation site on CHK2. We then generated site-specific point mutants of CHK2. Mutating T378 to Ala (T378A) largely reduced the O-GlcNAc signal compared with WT CHK2 and the T383A mutant control (Fig. 4o), indicating that T378 is the major CHK2 site carrying the O-GlcNAc modification. The potentially O-GlcNAcylated residue T378 and the surrounding amino acids are highly conserved among vertebrates (Fig. 4p), indicating that it serves an evolutionarily conserved role regulating the CHK2 protein. Taken together, these data indicate that CHK2 interacts with and can be O-GlcNAcylated by OGT in PKO cells.

### O-GlcNAcylation on T378 Stabilizes CHK2 and Promotes Hepatoma Cell Proliferation

To examine the effect of O-GlcNAcylation on CHK2 under the low-glucose conditions, Flag-tagged WT and T378A CHK2 were overexpressed with HA-tagged ubiquitin (HA-Ub) in PCK1-KO and parental PLC/PRF/5 cells. WT CHK2 ubiquitination was alleviated in PKO cells compared with parental cells, whereas the T378A mutation or OGT inhibitor ST045849 enhanced CHK2 ubiquitination (Fig. 5a). We then performed a series of cycloheximide-chase experiments to assess the half-life of these proteins. Endogenous CHK2 was more stable, with half-life of over 24h, in PLC/PRF/5 cell treated with PUGNAc, indicating CHK2 O-GlcNAcylation may enhance its stability (Extended Data Fig. 5a,b). In comparison with the parental cells, the CHK2 half-life was prolonged in PKO cells, but the T378A mutation or ST045849 treatment reduced CHK2 half-life from 24 h to 12 h (Fig. 5b-e). In addition, overexpression of WT PCK1 promoted CHK2 ubiquitination and degradation (Extended Data Fig. 5c-g). As expected, the G309R PCK1 mutation did not affect the half-life of CHK2 (Extended Data Fig. 5d-g). These results suggested that O-GlcNAc modification of T378 stabilizes CHK2 by preventing its ubiquitination and degradation in PCK1-deficient hepatoma cells.

**Fig. 5.**
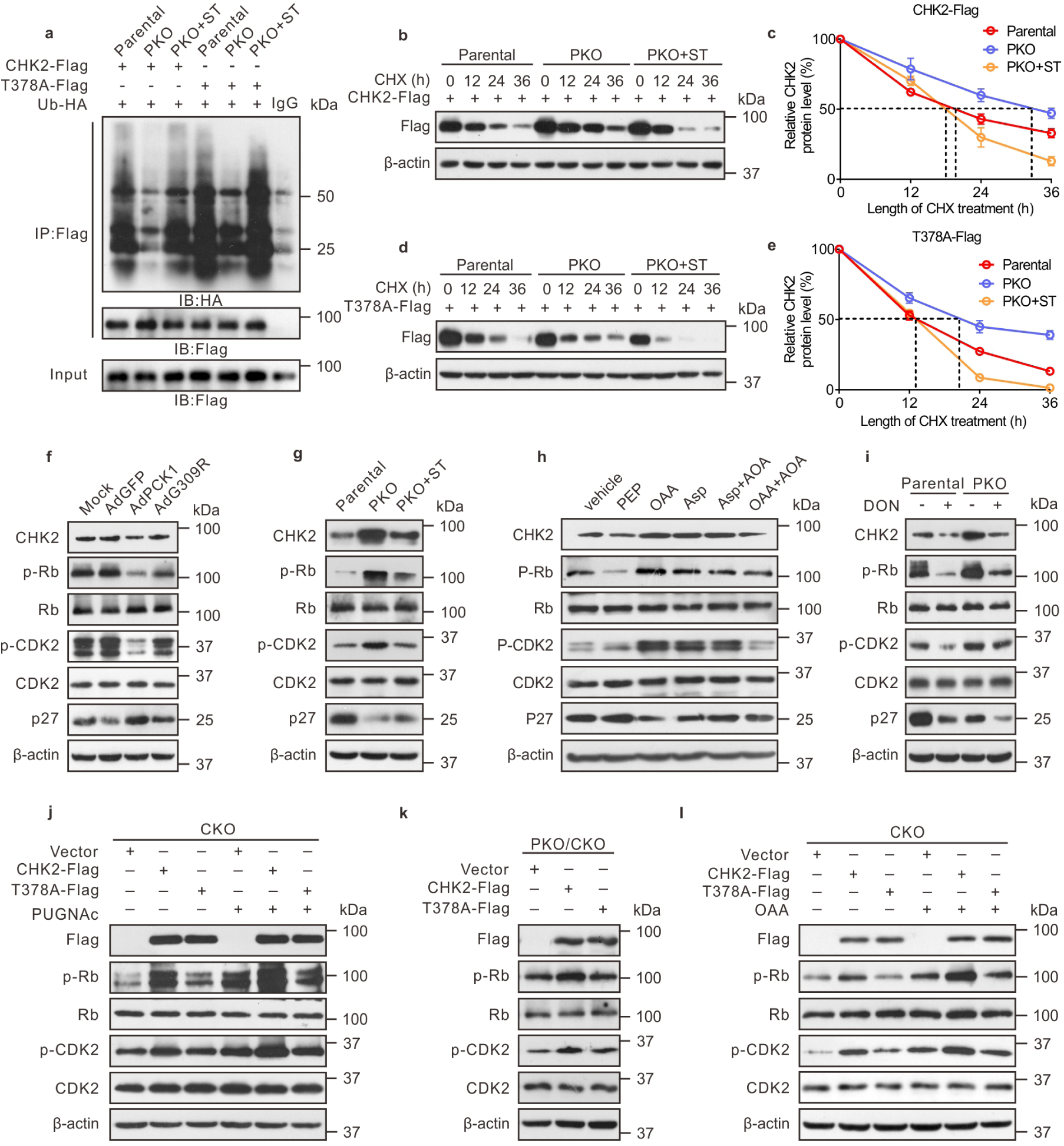
O-GlcNAcylation at T378 Stabilizes CHK2 and Activates Its Downstream Targets. **a**, CHK2 ubiquitination in PKO cells in the presence of HA-tagged ubiquitin (Ub-HA). **b**-**e**, Half-life and quantitative analysis of Flag-tagged WT CHK2 (**b**,**c**) and T378A mutant CHK2 (**d**,**e**) in PKO cells. Cells were treated with 40 μM cycloheximide (CHX) for the indicated time, and CHK2 levels was analyzed by immunoblotting. **f**-**i**, Representative immunoblots of CHK2, p-Rb, p-CDK2, and p27 expression in PCK1-OE cells (**f**), PKO cells (**g**,**i**), or SK-Hep1 cells following the indicated treatments (**h**). **j**,**l**, CKO cells (CHK2-knockout SK-Hep1 cells) were transfected with vectors CHK2-Flag or T378A-Flag, followed by treatment with 50 μM PUGNAc for 24 h (**j**) or 1 mM OAA for 12 h (**l**). Cells were lysed and analyzed by western blotting. **k**, PCK1/CHK2 double-knockout PLC/PRF/5 cells (PKO/CKO cells) were transfected with a CHK2-Flag or T378A-Flag expression vector, followed by Immunoblotting.

Given that CHK2 dimerization is essential for its activation ^29^, we next detected CHK2 dimerization in cells co-transfected with vectors encoding Flag-CHK2 and Myc-CHK2. Our results indicated that PKO cells displayed strengthened CHK2 dimer formation, whereas ST045849 treatment weakened this association (Extended Data Fig. 5h). A similar result was observed by crosslinking analysis (Extended Data Fig. 5i), suggesting O-GlcNAcylation of CHK2 may promote its dimerization. Interestingly, T378, the autophosphorylation site of CHK2, is located in the dimerization interface ^29^. We then performed dimeric CHK2 homology modeling, followed by molecular dynamic (MD) simulation. Our model disclosed that the O-GlcNAcylated residue T378 interacts with the amino acids VSLK of another CHK2 kinase domain. The acetylglucosamine group occupies a cavity which locates in the edge of interaction interface and forms three hydrogen bonds with the backbone of VSLK motif, thus might strengthen the stability of the CHK2 dimer (Extended Data Fig. 5j-l). Since dimerization promotes CHK2 activation and phosphorylates its downstream targets, such as retinoblastoma (Rb) in HCC ^30^, we subsequently checked the phosphorylation of CHK2 substrates and downstream signaling in response to PCK1 expression. Overexpressing WT PCK1 decreased the p-Rb and p-CDK2 levels, but increased the p27 levels (Fig. 5f). In contrast, PCK1 KO or OAA treatment reversed the regulatory effects of these molecules (Fig. 5g,h). Notably, the OGT inhibitor ST045849 or the GFAT1 inhibitor DON partially offset the regulatory effects mediated by PCK1 deficiency (Fig. 5g,i). These data indicated that O-GlcNAcylation promotes CHK2 dimerization and subsequently enhances downstream Rb phosphorylation.

To further test whether the loss of CHK2 O-GlcNAcylation affects its downstream signaling and hepatoma cell proliferation, we transiently overexpressed WT CHK2 or the T378A mutant in CHK2-KO cells. WT CHK2 restored the phosphorylation of Rb and CDK2 and promoted HCC proliferation, whereas the CHK2 T378A mutant failed to exert this stimulatory role on tumorigenesis (Fig. 5j and Extended Data Fig. 6a,b), indicating that T378 O-GlcNAcylation plays an essential role in CHK2 activation. The T378A point mutation (which eliminated the O-GlcNAc modification) decreased the capacity of CHK2 to phosphorylate Rb. In agreement, the OGA inhibitor PUGNAc enhanced the ability of WT CHK2 to phosphorylate Rb and CDK2, but not that of the CHK2 T378A mutant (Fig. 5j). Accordingly, PCK1 KO or OAA treatment promoted WT CHK2 activation (Fig. 5k,l). Finally, we explored whether PCK1 deficiency-induced malignancy may rely upon CHK2 O-GlcNAcylation. As expected, CHK2 depletion suppressed PKO cell proliferation, which was rescued by re-expressing WT CHK2, but not the T378 mutant (Extended Data Fig. 6c-f). These data suggest that CHK2 T378 O-GlcNAcylation conferred a growth advantage for the PKO cells. Collectively, these findings indicate that the O-GlcNAcylation of residue T378 stabilizes CHK2 and activates its downstream targets such as Rb, thus promoting PCK1-deficient hepatoma cell proliferation *in vitro*.

**Fig. 6.**
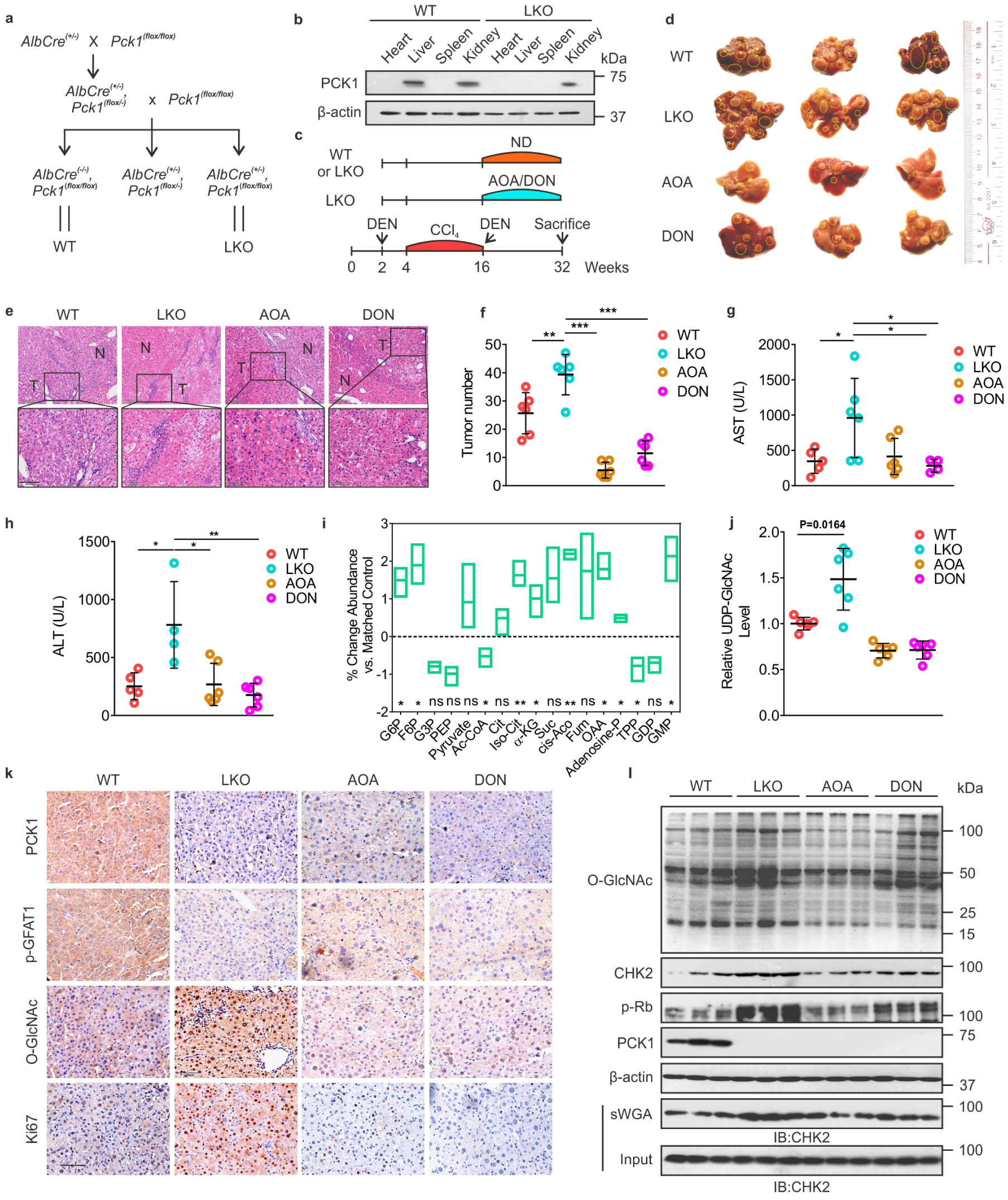
O-GlcNAcylation Promotes DEN/CCl_4_-Induced Hepatocellular Carcinogenesis in PCK1-Knockout Mice. **a**, Reproductive strategy for generating *AlbCre*^*(-/-)*^, *Pck1*^*(flox/flox)*^ (WT), and *AlbCre*^*(+/-)*^, *Pck1*^*(flox/flox)*^ (liver-specific knockout, LKO) mice. **b**, PCK1 protein expression in WT and LKO mouse organs involving the heart, liver, spleen, and kidney were confirmed by immunoblotting. **c**, Schematic representation of the experimental procedures used with WT and LKO mice. Mice were injected intraperitoneally with 75 mg/kg DEN or 4% CCl_4_ (every 3 days) as indicated, followed by combined administration of 5 mg/kg AOA or 1 mg/kg DON (twice per week) for 16 weeks. Control mice were provided a normal diet (ND). **d**-**f**, Gross appearances (**d**) and hematoxylin and eosin staining (**e**) of liver samples with tumors, and the numbers of tumor nodules (**f**). n = 6/group. The yellow dotted-line circles represent tumors. **g**,**h**, AST (**g**) and ALT (**h**) levels in mouse serum samples (n = 6/group). **i**, Glycolysis-metabolite profiles, derived from liver tumors of WT or LKO mice, were determined by performing LC-MS/MS metabolomics assays. **j**, Relative UDP-GlcNAc levels (n = 6/group). **k**,**l**, The indicated proteins in liver tumors were assessed by immunohistochemical labeling (**k**, scale bars: 100 μm) and western blotting (**l**).

### Targeting HBP-Meditated O-GlcNAcylation Suppresses DEN/CCl_4_-Induced Hepatocarcinogenesis *in Vivo*

Next, we used the DEN/CCl_4_-induced mouse model of liver cancer to further verify our results *in vivo*. Based on our *in vitro* data, we proposed that blocking the HBP with an inhibitor of GOT2 (AOA) or GFAT1 (DON) could suppress the growth of HCC by reducing O-GlcNAcylation. LKO mice (Fig. 6a,b) were generated as described previously ^31^. The mice were treated with DEN/CCl_4_ to induce hepatocarcinoma, which was followed by administering AOA or DON (twice a week) for 16 weeks (Fig 6c). The LKO mice exhibited accelerated liver tumorigenesis with increased tumor masses and nodules, and higher serum levels of aspartate aminotransferase (AST) and alanine aminotransferase (Fig. 6d-h). Our MS data showed that the OAA and UDP-GlcNAc levels were higher in the liver tumors of LKO mice (Fig. 6i,j), indicating that PCK1 deficiency promotes the HBP *in vivo*. In contrast, mice treated with AOA or DON exhibited slower tumor growth and a reduced number of tumor nodules, compared with those of untreated LKO mice (Fig. 6d-f). Furthermore, AOA or DON treatment also decreased UDP-GlcNAc and O-GlcNAcylation levels in LKO mice (Fig. 6j-l). These data suggested that hyper-O-GlcNAcylation conferred a growth advantage for tumor cells *in vivo*. Consistent with our *in vitro* data, the levels of total CHK2, O-GlcNAcylated CHK2, and p-Rb were significantly enhanced in the liver tumors of LKO mice, which was partially reversed by administering AOA or DON (Fig. 6l). In summary, these findings suggested that PCK1 depletion increases susceptibility to DEN/CCl_4_-induced carcinogenesis and promotes hepatocarcinogenesis via enhanced CHK2 O-GlcNAcylation; thus, blocking HBP-meditated O-GlcNAcylation suppresses HCC in LKO mice.

### PCK1 Deficiency Strengthens CHK2 O-GlcNAcylation in Primary Human HCC

Finally, we investigated PCK1 expression and global O-GlcNAcylation in 40 paired human primary HCC tissues and tumor-adjacent normal tissues. As shown by our immunohistochemistry (IHC) and immunoblot results, PCK1 was down-regulated in most HCC tissues (Fig. 7a-c and Extended Data Fig. 7), and deficient PCK1 expression was significantly associated with a larger tumor size and accelerated proliferation (Fig. 7a,d, Pearson correlation’s coefficient (r) = -0.3935, p = 0.0160; Supplemental Table 2). In addition, downregulated PCK1 expression was significantly associated with poor tumor differentiation and prognosis (Supplemental Table 2). Moreover, the global O-GlcNAcylation was significantly higher in HCC tissues than in adjacent normal tissues (Fig. 7b,e). Consistently, the p-AMPK and p-GFAT1 levels were reduced in HCC tissues (Fig. 7b). We also observed a negative correlation between PCK1 protein-expression levels and O-GlcNAcylation levels in HCC (Fig. 7f, r = -0.3565, p = 0.0240). In addition, the sWGA pull-down assay showed that enhanced CHK2 O-GlcNAcylation was associated with PCK1 down-regulation (Fig. 7g,h). Consistent with our *in vitro* data, we observed a strong negative correlation between p-Rb levels and PCK1 expression (Fig. 7i, r = -0.5347, p = 0.0270). In conclusion, this clinical validation supports the finding that PCK1 repression strengthens CHK2 O-GlcNAcylation and promotes tumor growth in human primary HCC.

**Fig. 7.**
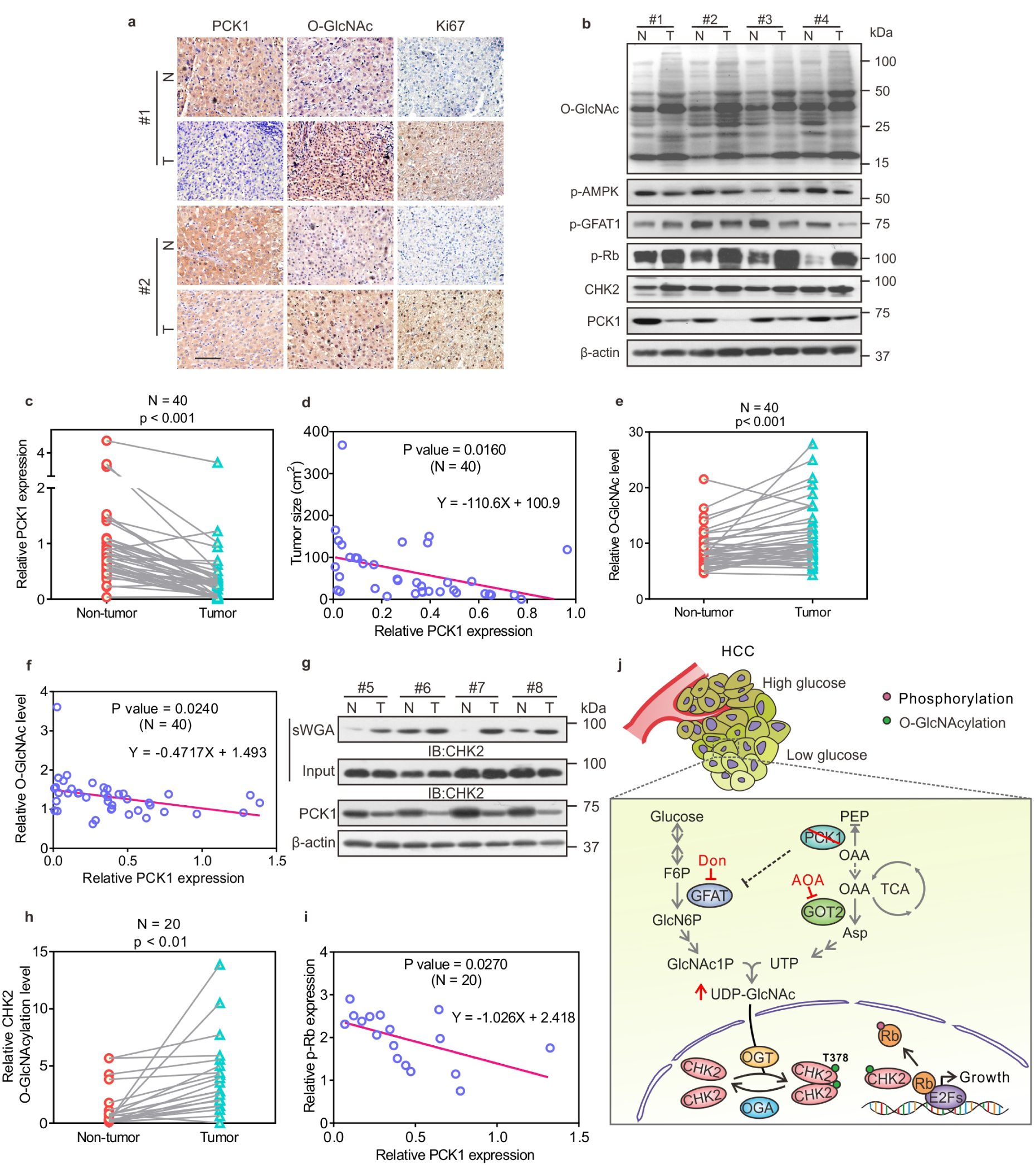
PCK1 Deficiency Strengthens Protein O-GlcNAcylation and Correlates with Human HCC Growth. **a**, IHC staining of PCK1, O-GlcNAcylation, and Ki67 in clinical HCC samples. Scale bars: 100 μm. **b**, Representative human HCC samples were indicated by immunoblots. **c**, Relative PCK1 protein-expression levels were compared between non-tumor tissues (NT) and tumors (T) from 40 patients with HCC (see also Figure S6). Relative protein-expression levels were normalized to those in NT samples. **d**, The correlation between HCC tumor sizes (n = 40) and PCK1 expression. **e**, Relative O-GlcNAc levels of proteins in samples from 40 patients (see also Figure S6). **f**, Correlation analysis between PCK1 expression and O-GlcNAc levels (n = 40). **g**,**h**, Analysis of CHK2 O-GlcNAcylation in HCC tumors by performing sWGA pull-down assays (**g**). CHK2 O-GlcNAcylation levels were quantified (**h**). **i**, Correlation analysis between PCK1 and p-Rb expression in tumor tissues from 20 patients with HCC. Data are represented as mean ± SD. P values were derived from Pearson’s correlation coefficient (r). **j**, Molecular Model for the role of PCK1 Deficiency in regulating CHK2 O-GlcNAcylation and HCC Growth upon low glucose. In a low-glucose microenvironment, PCK1 ablation promotes oxaloacetate accumulation and GFAT1 activation to increase UDP-GlcNAc synthesis through the hexosamine-biosynthesis pathway. Increased O-GlcNAc modification enhances Thr378 O-GlcNAcylation in CHK2, which leads to its dimerization and Rb phosphorylation, and HCC cell proliferation. Inhibitors AOA and DON suppress HCC growth, indicating a unique potential for targeting O-GlcNAc signaling in the treatment of HCC.

## DISCUSSION

Emerging evidence has demonstrated that protein O-GlcNAcylation plays key role in tumorigenesis. Increased glucose flux through the HBP elevates UDP-GlcNAc, which enhances cellular O-GlcNAcylation. However, cancer cells are frequently faced with limited nutrients due to an insufficient and inappropriate vascular supply and rapid nutrient consumption ^2^. Previous metabolomics data demonstrated that the glucose concentrations in tumor tissues are generally lower than those in non-transformed tissues ^32^. The mechanisms underlying tumor growth during periods of metabolic stress through enhanced HBP activity and O-GlcNAcylation have not been fully elucidated. It remains unknown whether gluconeogenesis contributes to maintaining HBP-mediated O-GlcNAcylation in cancer cells under low nutrient conditions ^4^. Here, we present the first evidence that deficiency of the gluconeogenic enzyme PCK1 promotes cellular O-GlcNAcylation and tumorigenesis in HCC (Fig. 7j). Moreover, we identify that CHK2 O-GlcNAcylation at T378 maintains its stability and oncogenic activity in hepatoma cells. Therefore, our study provides a link between PCK1 repression and hyper-O-GlcNAcylation underlying HCC oncogenesis.

Since the liver is the major site of gluconeogenesis during fasting, the role of gluconeogenesis in HCC has begun to draw more attention recently. During glucose starvation, cancer cells redistribute gluconeogenic intermediates to downstream pathways to facilitate their proliferation ^4^. PCK1, the rate-limiting enzyme of gluconeogenesis, is downregulated in HCC ^33,34,25^. PCK1 mediates not only gluconeogenesis, but also glyceroneogenesis and TCA cataplerosis ^35^. In this study, we found that PCK1 silencing promoted the HBP and UDP-GlcNAc biosynthesis, thus enhancing cellular O-GlcNAcylation under a low-glucose condition (5 mM glucose).

Interestingly, previous findings showed that the maximum rate of gluconeogenesis is approached at glucose concentrations under 5 mM, whereas high glucose levels inhibit gluconeogenesis ^36,37^. Consistent with this observation, we did not detect any significant change in O-GlcNAcylation levels under high-glucose conditions (25 mM glucose) upon PCK1 depletion or overexpression, indicating that PCK1 regulated HBP flux and protein O-GlcNAcylation, depending on glucose availability.

O-GlcNAcylation depends on OGT/OGA levels, and the HBP produces the donor substrate UDP-GlcNAc. Since PCK1 did not affect the expression levels of OGT and OGA, we studied the effect of PCK1 on HBP-mediated UDP-GlcNAc biosynthesis. As a nutrient sensor, UDP-GlcNAc levels are dependent on glucose, amino acid, fatty acid, and nucleotide availability ^38^. Here, we revealed a dual role for PCK1 in regulating UDP-GlcNAc biosynthesis through OAA accumulation and the AMPK-GFAT1 axis, under low-glucose conditions. On the one hand, the OAA levels decreased upon PCK1 overexpression in hepatoma cells, but accumulated in the liver tumors of LKO mice. LKO mice are unable to remove oxaloacetate from the TCA cycle ^39^. OAA is converted to Asp by GOT2, a key enzyme that plays a role in the TCA cycle and amino acid metabolism ^40^. Accordingly, amino acids synthesized from the TCA cycle (including Asp) were elevated in the blood of PCK1 KO mice ^41^.

Pharmacological or transcriptional inhibition of GOT2 suppressed hyper-O-GlcNAcylation induced by PCK1 deficiency or additional OAA treatment *in vitro*. Moreover, treatment with the GOT2 inhibitor, AOA reduced UDP-GlcNAc and O-GlcNAcylation levels in LKO mice, which may represent an important therapeutic perspective for HCC treatment. These findings implied that the cataplerotic function of PCK1 and the GOT2-mediated pathway are involved in regulating UDP-GlcNAc biosynthesis. Whether OAA-derived Asp is incorporated into UDP-GlcNAc via pyrimidine synthesis requires further study, based on stable isotope tracing.

On the other hand, our previous work showed that enforced PCK1 expression leads to energy reduction and activates AMPK upon glucose deprivation in HCC ^25^. GFAT1 is an AMPK substrate, so we hypothesized that PCK1 may suppress HBP-mediated O-GlcNAcylation through the AMPK-GFAT1 axis. In support of this hypothesis, we found that PCK1 regulated GFAT1 phosphorylation and O-GlcNAcylation levels in an AMPK-dependent manner, under glucose restrictions. As a rate-limiting enzyme of HBP, GFAT1 phosphorylation at Ser243 by AMPK diminishes its enzymatic activity ^42^, which decreases UDP-GlcNAc biosynthesis and O-GlcNAcylation levels. Interestingly, the AMPK-GFAT1 axis was also reported to regulate protein O-GlcNAcylation in angiogenesis and cardiac hypertrophy, indicating that this signal axis plays multiple physiological roles ^26,27^. In addition, AMPK directly phosphorylated residue Thr444 of OGT, the enzyme responsible for O-GlcNAcylation ^43^. Therefore, PCK1 may regulate the HBP and O-GlcNAcylation through different mechanisms.

It is known that O-GlcNAcylation is crucial for cell-cycle regulation and DNA-damage responses ^44^. Several proteins that regulate the growth and proliferation of tumor cells can be O-GlcNAcylated. For example, the G1/S checkpoint protein Rb is heavily O-GlcNAcylated during the G1 phase ^45^. By characterizing the role of O-GlcNAcylation upon PCK1 deficiency, we uncovered CHK2 as novel target of OGT. CHK2, a cell-cycle checkpoint kinase, play key roles in DNA-damage responses and cell-cycle progression. CHK2 is expressed in the nucleus in a subset of HCC and correlates with HCC progression ^30^. Several post-translational modifications, including phosphorylation, ubiquitination, and acetylation have been reported to be critical for CHK2 function ^46–48^. Herein, we identified Thr378 as a key O-GlcNAcylation site on CHK2 using LC-MS/MS. Importantly, the loss of O-GlcNAcylation by the T378A mutation increased CHK2 ubiquitination, thus promoted its degradation. Moreover, we found that O-GlcNAcylation promoted CHK2 dimerization and activation, therefore enhancing Rb phosphorylation in PCK1-deficient hepatoma cells. Further structural analysis of O-GlcNAcylated CHK2 may help us to understand how O-GlcNAcylation contributes to CHK2 activation.

Histopathology had revealed increased O-GlcNAcylation levels in HCC tumor tissues ^49^. Our findings not only provide an underlying mechanism whereby disrupted gluconeogenesis may activate the HBP and increase the availability of UDP-GlcNAc for O-GlcNAcylation under nutrient limitations, but also provides potential therapeutic targets for HCC. Preclinical evaluation of DON and AOA through the inhibition of glutamine metabolism has provided promising results for acute myeloid leukemia ^22^, high MYC-expressing atypical teratoid/rhabdoid tumors ^50^, and breast cancer ^51^. Here, we showed that both DON and AOA inhibited the growth of HCC *in vitro* and *in vivo*, largely by blocking HBP-mediated O-GlcNAcylation. These data suggest that DON and AOA inhibit cell growth through a novel mechanism and provide a strong rationale for further clinical drug development, particularly for PCK1-deficient HCC.

In summary, we uncovered a link between gluconeogenesis disruption and O-GlcNAcylation upon glucose deprivation in HCC. We demonstrated that PCK1 deficiency can promote HBP-mediated UDP-GlcNAc biosynthesis through OAA accumulation and the AMPK-GFAT1 axis. Moreover, the OGT-mediated O-GlcNAcylation of CHK2 on Thr378 stabilizes CHK2 and promote its oncogenic activity in HCC. The results of this study expands our understanding of PCK1 in hepatic carcinogenesis and indicates the potential of targeting HBP-mediated O-GlcNAcylation for HCC therapy.

## METHODS

### Cell Culture and Treatment

PLC/PRF/5, SK-Hep1, Huh7, MHCC-97H, and HEK293 cells were cultured in DMEM (HyClone, Logan, UT, USA) supplemented with 10% FBS (Gibco, Rockville, MD, USA), 100 U/ml penicillin, and 100 mg/ml streptomycin (HyClone) at 37°C in 5% CO_2_. For low-glucose treatment, cells were briefly washed with PBS (DINGGUO, BF-0011) and then maintained in glucose-free medium (Gibco, 11966025) supplemented with 10% FBS and glucose at various concentrations for 12 h. In addition, 3-MPA, Dex, ST, TG, AOA, DON, or metformin was added to the medium, as indicated.

### Animal Studies

*AlbCre*^(+/-)^, *Pck1*^(flox/flox)^ (LKO) mice were generated from crosses between *AlbCre*^*(+/-)*^ mice (Model Animal Research Center of Nanjing University, Nanjing, China) and *Pck1*^(flox/flox)^ mice with a C57BL/6 background (Mutant Mouse Resource & Research Centers, MMRRC:011950-UNC), as described previously ^31^, and *AlbCre*^(-/-)^, *Pck1*^(flox/flox)^ (WT) mice were used as a control (n = 6/group). HCC was induced in mice by combined treatment with DEN (75 mg/kg) and CCl_4_ (2 ml/kg, twice per week for 12 weeks), as described previously ^52^. At 16 weeks after DEN/CCl_4_ treatment, the LKO mice were administered an intraperitoneal injection of 5 mg/kg AOA-hemihydrochloride (Sigma) or 1 mg/kg DON (Sigma) twice a week for 16 weeks. At 32 weeks, the mice were sacrificed after fasting for 12 h, and liver tissues with tumors were collected for examination. Mouse serum ALT and AST were detected using an automatic biochemical analyzer (Catalyst One, IDEXX, USA).

For the orthotopic implantation model, BALB/c nude mice (n = 6/group) were randomly divided into 5 groups. For each nude mouse, AdGFP-, AdPCK1-, AdG309R-, or mock-infected MHCC97H cells (1 × 10^5^) were suspended in a 50-µl PBS/Matrigel (356234, BD Biosciences) mixture (1:1 ratio, v/v) and then implanted into the left liver lobe. At 14 days after injection, the Dex group was intraperitoneally injected with 5 mg/kg Dex per day for 14 days. At 4 weeks after implantation, all mice were sacrificed after fasting for 12 h. All animal procedures were performed according to protocols approved by the Institutional Animal Care and Use Committee at the Laboratory Animal Center of Chongqing Medical University. All procedures were also approved by the Research Ethics Committee of Chongqing Medical University (reference number: 2017012).

### Clinical Specimens

HCC samples and paired, adjacent normal liver tissues were obtained from the Second Affiliated Hospital of Chongqing Medical University between 2015 and 2018, with approval from the Institutional Review Board of Chongqing Medical University. Clinical samples were collected from patients after obtaining informed consent in accordance with a protocol approved by the Second Affiliated Hospital of Chongqing Medical University (Chongqing, China).

### Plasmid Constructs

For recombinant plasmid construction, DNA fragments (PCK1 NM_002591, CHEK2 NM_007194.4, ΔN truncated CHK2 69-543aa, ΔC truncated CHK2 1-221aa, and OGT NM_181672.2) were amplified by PCR and separately cloned into the pSEB-3Flag, pBudCE4.1-3HA, or pBudCE4.1-5Myc vector. PCK1 G309R, CHK2 T378A, and CHK2 T383A were constructed by overlapping PCR. To construct plasmids expressing shRNAs, two pairs of oligonucleotides encoding shRNAs targeting OGT, OGA, AMPK, or GOT2 (Supplemental Table 3) were cloned into the pLL3.7 vector (from Prof. Bing Sun, Shanghai Institute of Biochemistry and Cell Biology, Chinese Academy of Sciences, China).

### Adenovirus Production

The full-length cDNA fragment of PCK1 (NM_002591) or G309R (PCK1 mutation 925G>A) was inserted into the pAdTrack-TO4 vector (from Dr. Tong-Chuan He, University of Chicago, USA). Recombinant adenoviral, AdPCK1 and AdG309R, were generated using the AdEasy system as described previously ^25^. The adenoviral AdGFP, expressing only GFP, was used as a control.

### CRISPR/Cas9-mediated Knockout Cells

PCK1- or CHEK2-knockout cells (PKO or CKO cells) were established using the CRISPR-Cas9 system (from Prof. Ding Xue, the School of Life Sciences, Tsinghua University, Beijing, China), as described previously ^25^. Single-cell HCC clones stably expressing single guide RNA (sgRNA) sequences were propagated and validated by immunoblotting and DNA sequencing. The sequences of all oligonucleotides used to generate the knockout cell lines are listed in Supplemental Table 3.

### Cell Lysis and Western Blotting

Protein lysates from cells or liver samples were extracted using cell lysis buffer (Beyotime Biotechnology, Jiangsu, China) containing 1 mM phenylmethanesulfonyl fluoride (Beyotime). Equal volumes of protein samples were separated by 10% SDS/PAGE and electro-transferred to PVDF membranes (Millipore, Billerica, MA, USA). The immunoblots were probed with the indicated antibodies. Proteins bands were visualized with Chemiluminescent Substrate (Bio-Rad, USA) and exposed to CareStream Film (Kodak, USA). Quantification of bands in western blots was performed using Image-Pro Plus software.

### RNA Extraction, Reverse Transcription PCR, and Quantitative Real-Time PCR

Total RNA was extracted from cells using the TRIzol(tm) reagent (Invitrogen, Carlsbad, CA, USA) and reverse-transcribed into cDNA using the PrimeScript(tm) RT Reagent Kit (RR047A, TaKaRa, Japan). Quantitative real-time PCR analysis of target mRNA-expression levels was achieved using SYBR Green qPCR Master Mix (Bio-Rad, Hercules, CA, USA) and the CFX Connect Real-time PCR Detection System (Bio-Rad). The specific primers used are shown in Supplemental Table 3. All samples were analyzed in triplicate for each experiment. The relative gene-expression levels were normalized to β-actin and calculated using the 2^–ΔΔCt^ method.

### Cell-Proliferation Assay

Cells were re-seeded in 96-well plates at a density of 1 × 10^3^ cells/well and cultured in low-glucose medium containing 5 mM glucose for 5 days. The plates were scanned and phase-contrast images were acquired in real time every 24 h post-treatment.

Quantified time-lapse curves were generated using IncuCyte ZOOM software (ESCO, Changqi South Street, Singapore). For clone formation capacity, cells were seeded into 6-well plate at 800/well. After 7-10 days on 5 mM glucose condition, the cells were fixed with paraformaldehyde and then incubated in crystal violet. The clones were photographed and the number of clones was counted by Image-Pro Plus software, version 6.0 (Media Cybernetics, Inc., Bethesda, MD, USA).

### Metabolites Detection and Analysis

Cells were washed twice with pre-cooled physiological saline, and metabolites were extracted with 400 μL cold methanol/acetonitrile (1:1, v/v) to remove the protein. The mixture was centrifuged for 20 min (14,000 × *g*, 4°C). The supernatant was dried in a vacuum centrifuge. For LC-MS analysis, the samples were re-dissolved in 100 μL acetonitrile/water (1:1, v/v) solvent. Mouse liver tumor tissues (60 mg) were extracted with 1 mL cold 90% methanol. The lysates were homogenized twice using an MP homogenizer (24 × 2, 6.0 M/S, 60 s). The homogenates were sonicated on ice. After centrifugation at 14,000 × *g* for 20 min, the supernatant was dried and re-dissolved in 100 μL acetonitrile/water (1:1, v/v) solvent.

Untargeted metabolomics profiling of PCK1-overexpressing cells was performed using ultra-high-performance liquid chromatography (Agilent 1290 Infinity LC) coupled to with quadrupole time-of-flight mass spectrometry (UHPLC-QTOF/MS) at Shanghai Applied Protein Technology Co., Ltd ^53^. UDP-GlcNAc levels were quantified by performing targeted LC-MS/MS analysis (ACQUITY UPLC I CLASS, Xevo G2-S QTof). TCA-derived metabolites were detected by UHPLC, using an Agilent 1290 Infinity LC column coupled to 5500 QTRAP system (AB SCIEX) at Shanghai Applied Protein Technology Co., Ltd.

### Immunoprecipitation

PKO or SK-Hep1 cells were transfected for 48 h with a fusion vector expressing Flag-tagged fusion protein (OGT-Flag, CHK2-Flag, T378A-Flag, or T383A-Flag). Cells were treated with 50 μm PUGNAc for 12 h and resuspended in lysis buffer (50 mM Tris HCl, pH 7.4, 150 mM NaCl, 1 mM EDTA, and 1% Triton X-100) containing protease (Roche) and phosphatase (Beyotime Biotechnology) inhibitor cocktails. Pre-cleared lysates were incubated overnight with an anti-FLAG M2 affinity gel (Sigma, A2220) at 4°C.

For co-IP analysis, HEK293 cells were co-transfected with vectors expressing OGT-HA and either CHK2-Flag, ΔC-Flag, or ΔN-Flag. PKO cells were co-transfected with the CHK2-Flag and CHK2-Myc vectors. Pre-cleared cell lysates were incubated overnight with an anti-Flag or anti-HA antibody, and coupled with 40 μl protein A/G agarose beads. Immunoprecipitated complexes were eluted and subjected to immunoblotting using the indicated antibodies.

### sWGA Pull-Down Assay

Hepatic cells and liver tissues were lysed in a buffer containing 125 mM NaCl, 50 mM Tris (pH 7.4), 5 mM EDTA, and 0.1% NP-40. The lysate was denatured in glycoprotein-denaturing buffer and digested with PNGase (NEB P0704S) to remove N-linked glycoproteins. Pre-cleared lysates were incubated overnight with sWGA-conjugated agarose beads (Vector Laboratories, Burlingame, CA). Precipitated complexes were eluted and immunoblotted with the indicated antibodies.

### IP-MS assay

PKO cells transfected with Flag-tagged OGT before cultured in 5 mM glucose were collected and incubated overnight at 4°C with ANTI-FLAG M2 affinity gel (Sigma, A2220). Immunoprecipitated complexes were eluted and stained with Coomassie blue. Excised protein gel bands were sent to Shanghai Applied Protein Technology Co., Ltd. to identify OGT-binding proteins. After performing the reduction reaction and alkylation treatment, the samples were incubated with trypsin (mass ratio, 1:50). The tryptic products were desalted, lyophilized, and re-dissolved in 0.1% formic acid solution. MS analysis was performed using a Q Exactive MS instrument (Thermo Fisher) coupled to an Easy-nLC 1000 LC instrument (Thermo Fisher) in Shanghai Applied Protein Technology Co., Ltd. The raw MS data were analyzed using Mascot 2.2 software to identify immunoprecipitating proteins. Further, pathway-enrichment analysis was performed using the KEGG database (http://kobas.cbi.pku.edu.cn/), p values < 0.05.

### Immunofluorescence Assays

CHK2 and OGT staining was performed with anti-Flag (mouse 1:200) and anti-HA (rabbit 1:800) antibodies, using PKO cells co-transfected with CHK2-Flag and OGT-HA vectors. Specific signals were visualized using secondary antibodies (goat anti-rabbit IgG/TRITC or goat anti-mouse IgG/FITC). For nuclear staining, cells were treated with 1 μg/mL DAPI (10236276001, Roche Diagnostics GmbH, Mannheim, Germany). Stained sections were detected by a laser-scanning confocal microscope (Leica TCS SP8, Solms, Germany).

### CHK2 O-GlcNAcylation Site Mapping

MS was performed to identify CHK2 O-GlcNAcylation sites, as described previously ^54^. Briefly, CHK2 was immunoprecipitated from PKO cells transfected with CHK2-Flag and subjected to 10% SDS-PAGE for Coomassie blue staining. The band corresponding to CHK2 was excised from the gel and digested overnight with trypsin. Online LC-MS/MS system consisting of an Easy-nLC system and an Orbitrap Fusion Lumos Tribrid MS instrument (Thermo Scientific, Germany) equipped with a nanoelectrospray ion source were performed. The data from the Raw MS files were analyzed against the UniProt database with MaxQuant software (version 1.5.2.8). The first-search peptide tolerance was 20 ppm and the main-search peptide tolerance was 6 ppm. The MS/MS tolerance was 0.02 Da. The peptide-spectrum match and the false-discovery level was set to 1%. Matches between runs were determined and the minimum score for modified peptides was set to 40.

### CHK2 Protein-Stability Assay

PCK1-OE SK-Hep1 cells or PKO cells transfected with CHK2 WT or mutant plasmids were treated with 40 μM cycloheximide and harvested after 0, 12, 24, or 36 h. The cells were lysed, and CHK2 was detected using an anti-Flag antibody (Sigma).

### CHK2-Oligomerization Assay

The PKO cells were incubated with HEPES buffer (40 mM HEPES pH 7.5, 150 mM NaCl, 0.1% NP-40) supplemented with protease (Roche) and phosphatase (Beyotime) inhibitor cocktails for 15 min and centrifuged at 14,000 × *g* for 20 min at 4°C. Whole-cell extracts were cross-linked with 0.025% glutaraldehyde for 5 min at 37°C and then quenched with 50 mM Tris-HCl (pH 8.0). The samples were then separated on a 4–20% precast gel and analyzed by western blotting using the indicated antibodies.

### Molecular Dynamic Simulation

Missing loops and side chains of the dimeric CHK2 structure were complemented using SWISS-model with a previously reported CHK2 dimer structure as template (PDB code: 3I6U). The O-GlcNAcylation was added onto the predicted structure using PyMOL 2.3 software. Molecular dynamic simulation was conducted using GROMACS 2019 package ^55^. Topologies were generated by ACPYPE with Amber99sb force field parameters ^56^. After an initial energy minimization, 10 ns of unrestrained NPT MD simulation was performed under constant temperature of 300 K and constant pressure of 1 atm.

### Immunohistochemical (IHC) Assay

Human or mouse liver tissues were fixed in 4% paraformaldehyde for at least 24 h and embedded in paraffin, according to standard procedures. The sections were incubated overnight at 4°C with primary antibodies against PCK1 (1:500), p-GFAT (1:200), O-GlcNAc (1:100), or Ki67 (1:100). Subsequently, the slides were incubated with secondary anti-rabbit or anti-mouse IgG (ZSGB-BIO, Beijing, China) and visualized using 3, 3′-diaminobenzidine (ZSGB-BIO). The stained slides were scanned with a Pannoramic Scan 250 Flash or MIDI system, and images were acquired using Pannoramic Viewer software, version 1.15.2 (3DHistech, Budapest, Hungary). The mean staining intensity was analyzed using Image-Pro Plus software.

### Quantification and Statistical Analysis

Integral optical density (IOD) values were measured using Image-Pro Plus software (version 6.0) to determine the intensity of protein expression. The IOD was calculated as the mean density × the area. The relative protein-expression levels were normalized to an internal reference control, such as β-actin. All statistical analyses were performed using GraphPad Prism 7 (GraphPad Software Inc.). Data are represented as mean ± SD. Student’s t-test was used to compare two groups.

One-way ANOVA followed by Tukey’s test was used to compare more than two groups. Pearson’ correlation coefficient (r) was used to test linear correlations. Statistical significance was defined as p values < 0.05. *P < 0.05, **P < 0.01, ***P < 0.001.

## ACKNOWLEDGMENTS

We would like to thank Dr. T.-C He (University of Chicago, USA) for providing the pAdEasy system and critical reading of the manuscript. We are grateful to Prof. Ding Xue (Tsinghua University, China) for supplying the CRISPR/Cas9 system. We also thank Prof. Bing Sun (Shanghai Institute of Biochemistry and Cell Biology, China) for providing the pLL3.7 vector. We thank Zhimin Lu (Zhejiang University, China) for suggestions and critical reading of the manuscript. This work was supported by the China National Natural Science Foundation (grant no. 81872270), the Natural Science Foundation Project of Chongqing (cstc2018jcyjAX0254, cstc2019jcyj-msxmX0587), the Major National S&T program (2017ZX10202203-004), and the Science and Technology Research Program of Chongqing Municipal Education Commission (KJQN201900429).

## AUTHOR CONTRIBUTIONS

NT, AH, and KW conceived and designed the study. J Xiang, CC, RL and DG performed most experiments and analyzed the data. LC and WZ performed CHK2 O-GlcNAcylation site mapping. HD and LT helped with data analysis. LL generated CHK2 mutants. QG, XP, and J Xia assisted with mice experiments. LH performed molecular dynamic simulation. J Xiang, KW, and NT wrote the manuscript with all authors providing feedback.

## DECLARATION OF INTERESTS

The authors declare no competing interests.

## Notes

### Competing Interest Statement

The authors have declared no competing interest.

