## Extended Data Fig for "Gluconeogenic Enzyme PCK1 Deficiency Is Critical for CHK2 O-GlcNAcylation and Hepatocellular Carcinoma Growth upon Glucose Deprivation"

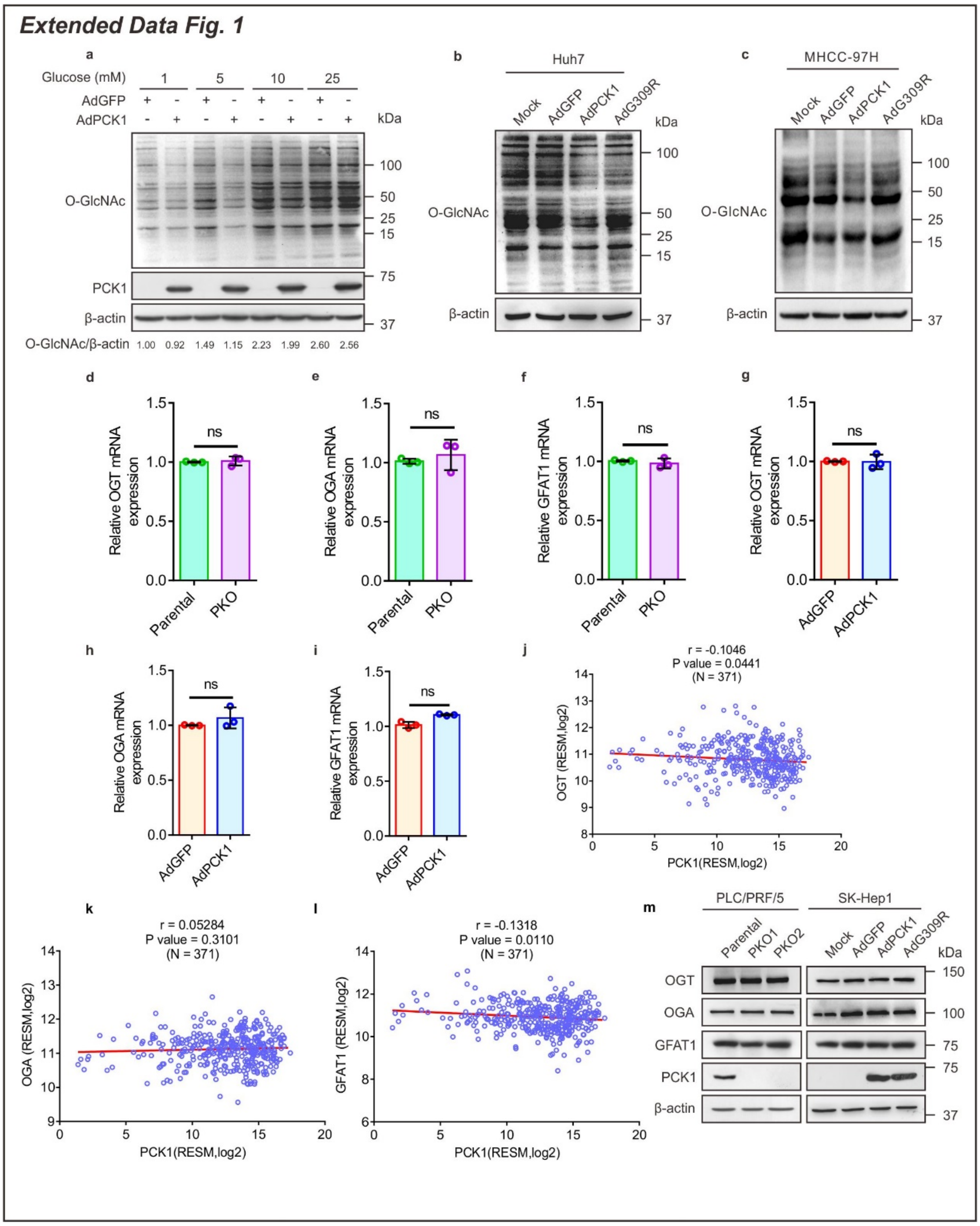


**Extended Data Fig. 1 The Expression of OGT, OGA and GFAT Are Not Regulated by PCK1. a**, Global O-GlcNAcylation levels in PCK1-expressing SK-Hep1 cells cultured in medium containing different concentrations of glucose for 12 h. Densitometric analysis was shown as indicated. **b**,**c**, Protein O-GlcNAcylation in Huh7 (**b**) and MHCC-97H cells (**c**) overexpressing PCK1 or

G309R, and then cultured with 5mM glucose medium for 12 h. **d**-**i**, OGT, OGA and GFAT1 mRNA expression in PKO cells (**d**-**f**) and PCK1-OE cells in SK-Hep1 (**g**-**i**) at 5 mM glucose. **j**-**l**, Correlation of PCK1 and OGT (**j**), OGA (**k**) or GFAT1 (**l**) mRNA expression in HCC patient samples from TCGA database. To compare the correlation of gene expression levels, we analyzed the RNA-sequencing data by Expectation Maximization (RSEM), two genes were normalized to transcript abundance of genes. **m**, The protein expression levels of OGT, OGA and GFAT1 in PKO cells (left) and SK-Hep1 cells overexpressing PCK1 or G309R (right) under low-glucose concentration (5mM). Data are represented mean ± SD (n ≥ 3), ns, not significant.

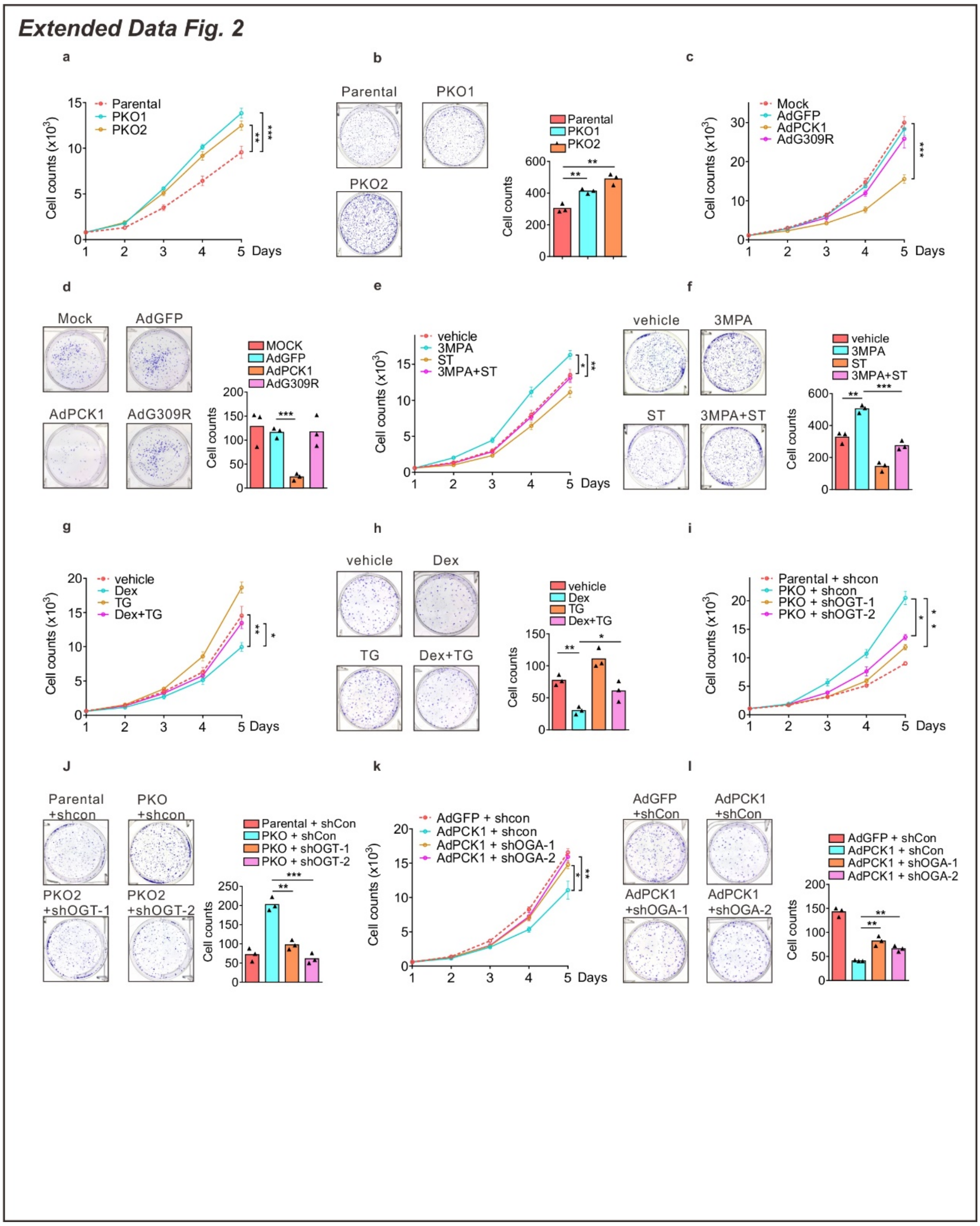


**Extended Data Fig. 2 PCK1 Inhibits Hepatoma Cells Proliferation via The HBP. a-d**, The growth curve and colony formation capacity of PKO cells (**a**,**b**) and PCK1-OE cells (**c**,**d**). **e**,**f**, SK-Hep1 cells were treated with 3-MAP (3-Mercaptopicolinic acid, PCK1 inhibitor, 200 μM) or ST (ST045849, OGT inhibitor, 100 μM) as indicated. **g**,**h**, PLC/PRF/5 cells were treated with TG

(Thiamet G, OGA inhibitor, 25 μM) or Dex (Dexamethasone, 10 μM) as indicated (**d**). **i**-**l**, PKO cells were transfected with OGT shRNA1/2 plasmid (**i**,**j**) and PCK1-OE cells were transfected with OGA shRNA1/2 plasmid (**k**,**l**), respectively. Values are shown as indicated (n≥3), $*p < 0.05$, $**p < 0.01$, $***p < 0.001$, one-way ANOVA followed by the Tukey's test.

*Extended Data Fig. 3*

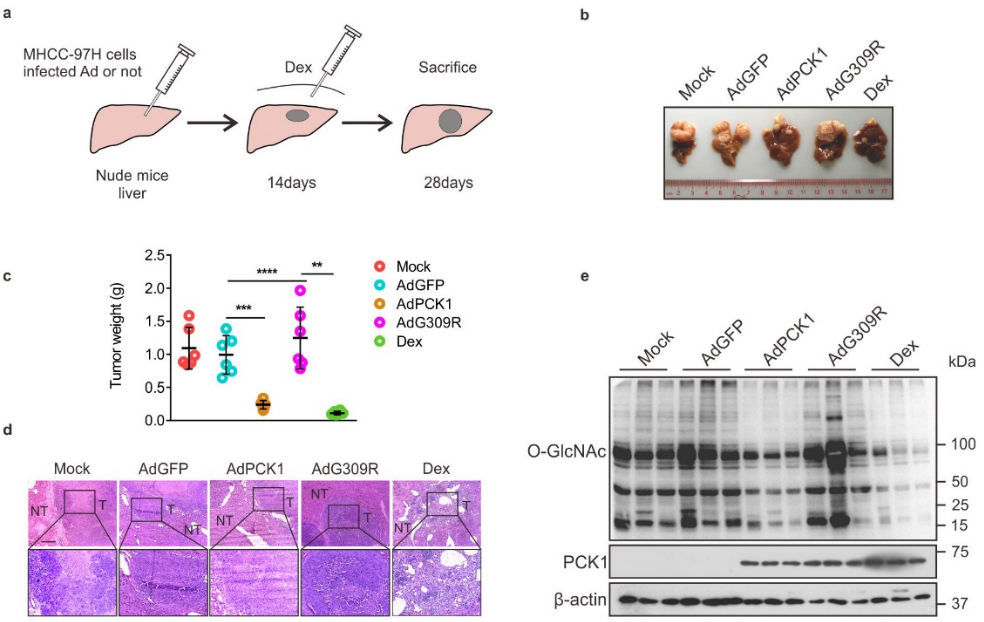


**Extended Data Fig.3 Overexpression of PCK1 Inhibits HCC Growth in *Vivo.*** **a**,**b**, An orthotopic implantation tumor model (**a**) established with MHCC-97H cells. Cells were infected with AdGFP, AdPCK1, or AdG309R, and implanted into the left liver lobes of nude mice. At 14 days after tumor formation, mice were administrated with 5 mg/kg Dex per day for 14 days. n =

6/group. **b**, Representative gross appearance of orthotopic implantation tumors. **c**,**d**, Tumor weight (**c**) and H&E staining analysis (**d**), Scale bars: 100 μm. **e**, Protein expression of PCK1 and total O-GlcNAcylation in liver tumors.

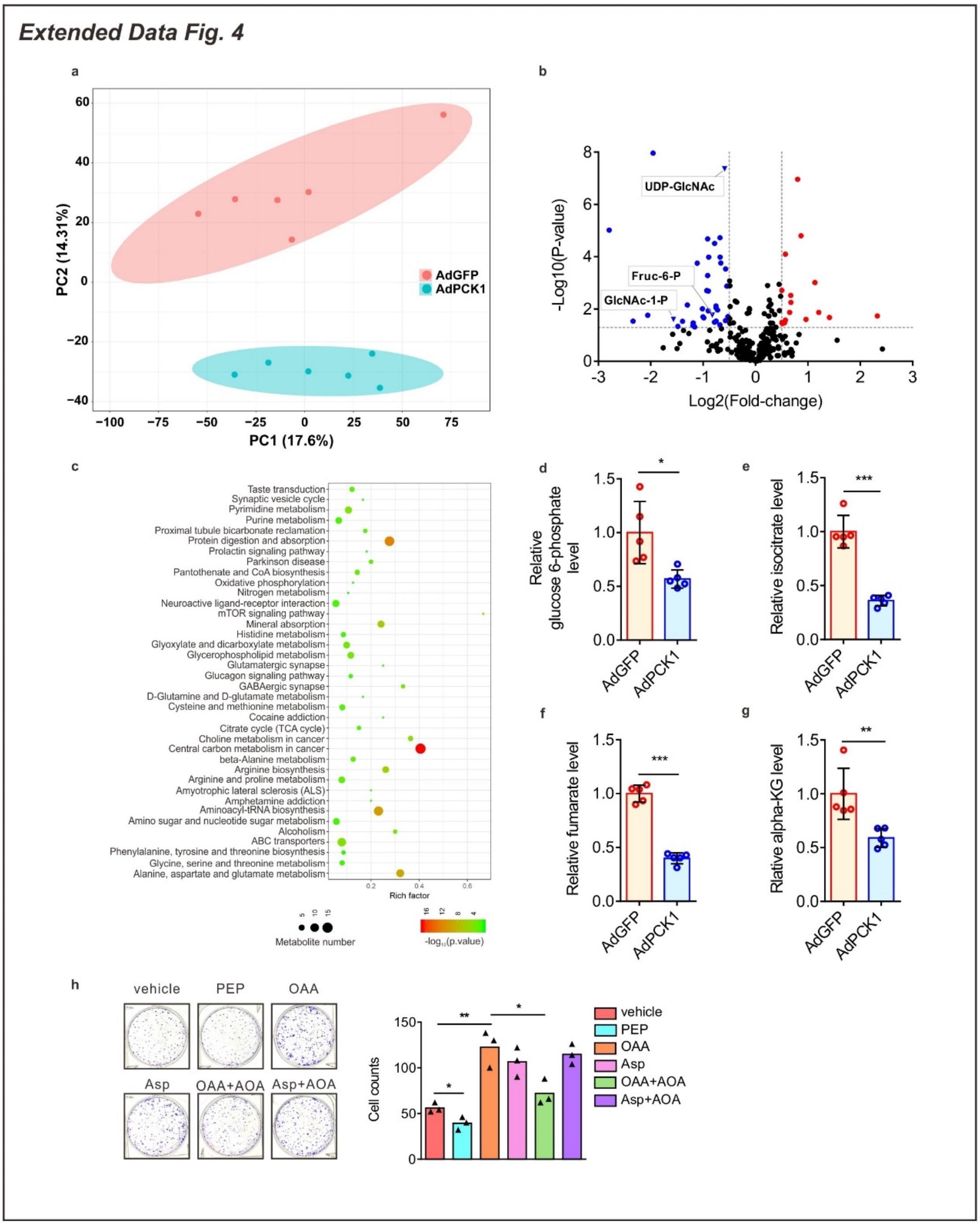


**Extended Data Fig. 4 Overexpression of PCK1 Inhibits Pyrimidine Metabolism. a-c**, Principal component analysis (**a**), volcano plot of fold-change including HBP metabolites (**b**), and pathway-enrichment analysis (**c**) of metabolite profiles were obtained using a metabolomics assay in SK-Hep1 cells infected with AdCPK1 or AdGFP and cultured in medium containing 5 mM of glucose for 12 h. **d-g**, LC-MS analysis of intermediate metabolites of glycolytic pathway and TCA cycle. **h**, Colony formation capacity of SK-Hep1 cells treated with 1 mM PEP, 1 mM OAA, 1 mM Asp or 20 μM AOA for 10 days. Values are shown as indicated, *$p < 0.05$, **$p < 0.01$, ***$p < 0.001$.

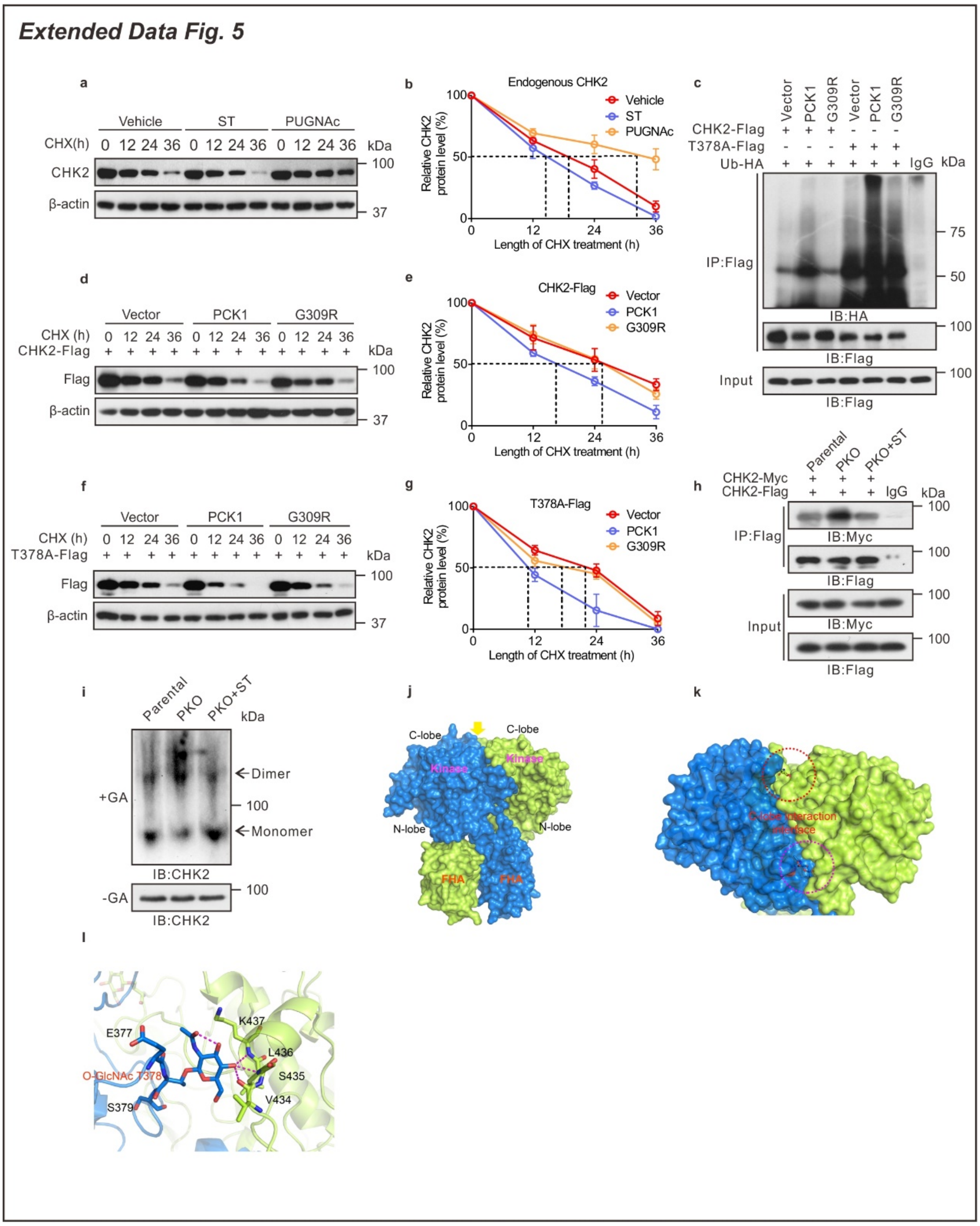


**Extended Data Fig. 5 PCK1 Promotes CHK2 Degradation and Reduces the Formation of CHK2 Dimer. a**,**b**, Half-life of endogenous CHK2 (**a**) in PLC/PRF/5 treated with PUGNAc or ST, CHK2 level was analyzed by using densitometric software (**b**). **c**, CHK2 ubiquitination in PCK1-OE cells in the presence of HA-tagged ubiquitin (Ub-HA). **d-g**, Half-life of CHK2-Flag (**d**) and

T378A-Flag (**f**) in PCK1-OE cells, densitometric analyses are shown in **e**,**g**. **h**, The co-IP of CHK2-Myc and CHK2-Flag was determined by anti-Flag antibody. **i**, The oligomer states of CHK2 in PKO cells. Cell lysis were treated with glutaraldehyde (GA) and probed with an anti-CHK2 antibody. **j**-**l**, Structural modeling of the dimeric CHK2 structure with O-GlcNAcylated T378. Dimeric CHK2 structures are covered with surface (**j**). Interaction interface of the C-lobe of the kinase domain (**k**). O-GlcNAcylated T378s are shown as sticks in the circles. Interactions between O-GlcNAcylated T378 and VSLK motif (**l**). E377, O-GlcNAcylated T378, and S379 are shown as marine blue sticks. The amino acids VSLK of the C-lobe of kinase domain are shown as light green sticks. Red dashed lines indicate hydrogen bonds.

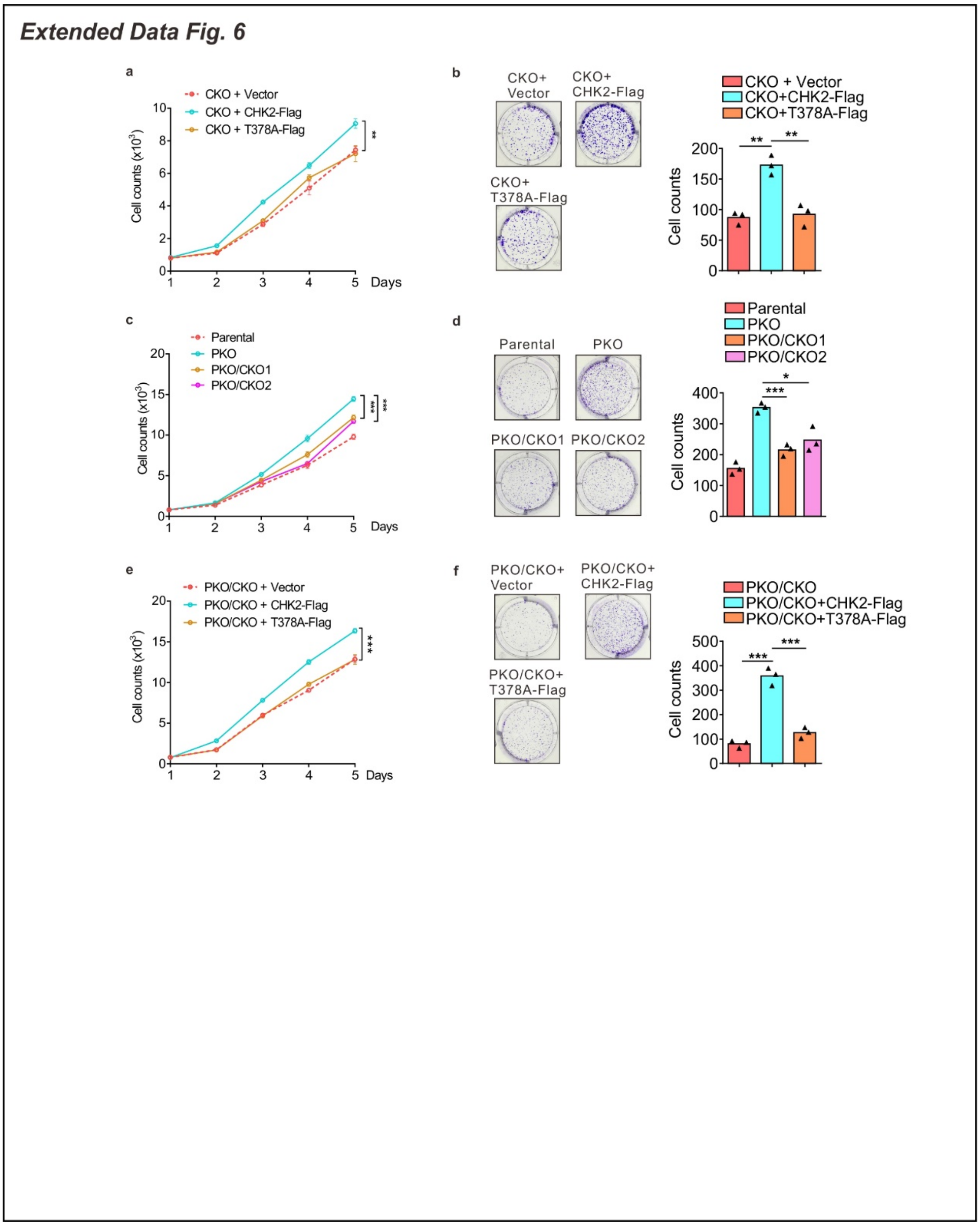


**Extended Data Fig. 6 CHK2 O-GlcNAcylation Promotes HCC Proliferation. a-f**, Cell proliferation and Colony formation analysis in CKO cells rescued by CHK2-Flag or T378A-Flag (**a**,**b**), and PKO/CKO cells (**c**,**d**) or rescue assay as indicated (**e**,**f**). Values are shown as indicated (n ≥ 3), *p < 0.05, **p < 0.01, ***p < 0.001, one-way ANOVA followed by the Tukey's test (more than two groups).

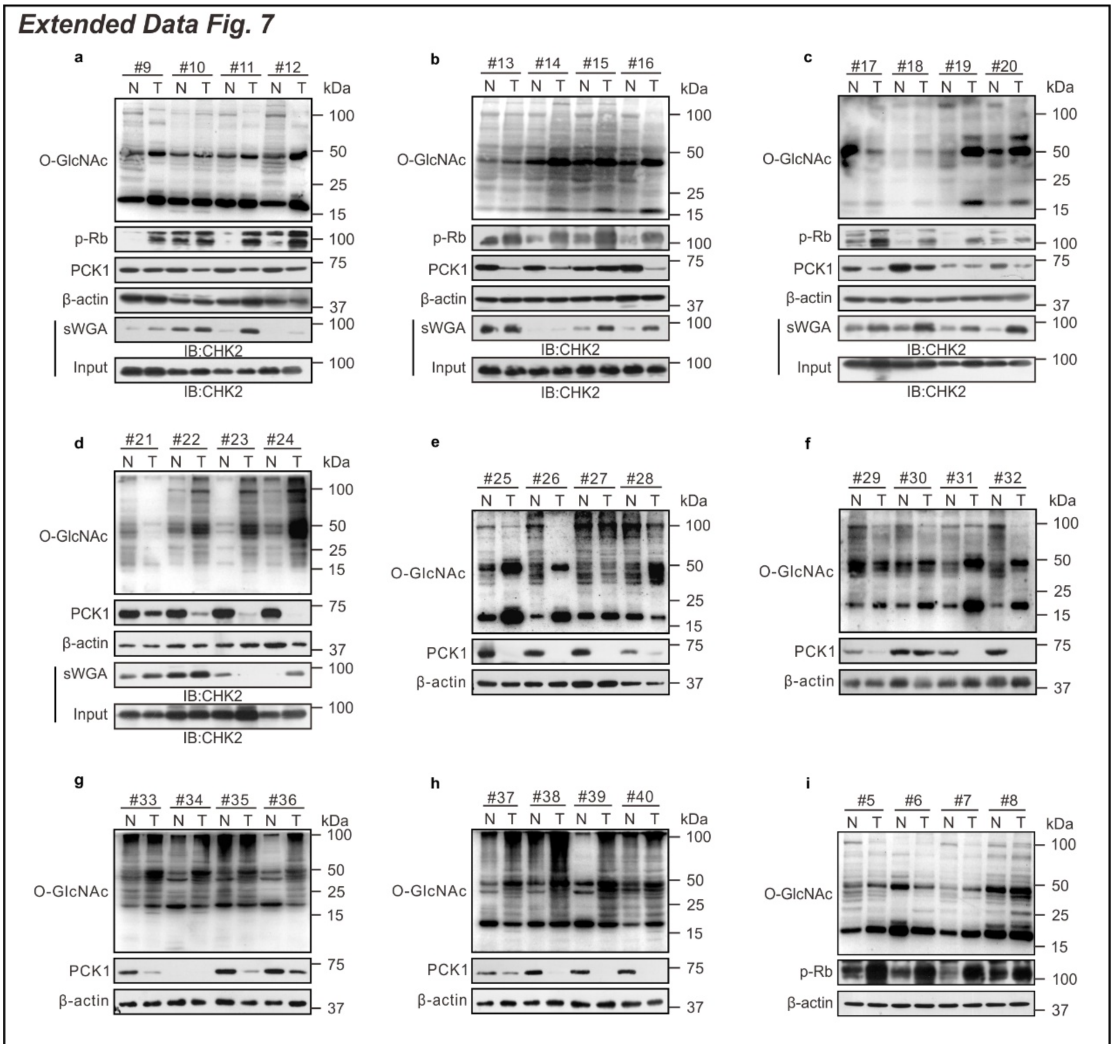


**Extended Data Fig. 7 Downregulation of PCK1 but Upregulation of O-GlcNAc in 40 Cases of HCC. a-i**, Protein expression of PCK1, global O-GlcNAcylation, p-Rb, and CHK2 O-GlcNAcylation in human HCC (See also **Fig. 7d**,**g**).
